# Comparative profiling of the synaptic proteome from Alzheimer’s disease patients with focus on the APOE genotype

**DOI:** 10.1101/631556

**Authors:** Raphael Hesse, Maica Llavero Hurtado, Rosemary J. Jackson, Samantha L. Eaton, Abigail G Herrmann, Marti Colom-Cadena, Declan King, Jamie Rose, Jane Tulloch, Chris-Anne McKenzie, Colin Smith, Christopher Henstridge, Douglas Lamont, Thomas M. Wishart, Tara L. Spires-Jones

**Author notes:** Corresponding authors (equal contributions) Contact: Tara Spires-Jones, UK Dementia Research Institute, Centre for Discovery Brain Sciences, The University of Edinburgh, 1 George Square, Edinburgh, EH8 9JZ, Scotland, UK, Thomas Wishart, Royal (Dick) School of Veterinary Studies, College of Medicine and Veterinary Medicine, University of Edinburgh, Easter Bush, Midlothian, EH25 9RG, SCOTLAND, UK. These authors contributed equally (Raphael Hesse, Maica Llavero Hurtado, Rosemary J. Jackson, Samantha L. Eaton).

## Abstract

Degeneration of synapses in Alzheimer’s disease (AD) strongly correlates with cognitive decline, and synaptic pathology contributes to disease pathophysiology. We recently discovered that the strongest genetic risk factor for sporadic AD, apolipoprotein E epsilon 4 (*APOE*4), exacerbates synapse loss and synaptic accumulation of oligomeric amyloid beta in human AD brain. To begin to understand the molecular cascades involved in synapse loss in AD and how this is mediated by *APOE*, and to generate a resource of knowledge of changes in the synaptic proteome in AD, we conducted a proteomic screen and systematic *in-silico* analysis of synaptoneurosome preparations from temporal and occipital cortices of human AD and control subjects with known *APOE* gene status. Our analysis identified over 5,500 proteins in human synaptoneurosomes and highlighted disease, brain region, and APOE-associated changes in multiple molecular pathways including a decreased abundance in AD of proteins important for synaptic and mitochondrial function and an increased abundance of proteins involved in neuroimmune interactions and intracellular signaling.

**Highlights:**

- Proteomic analysis of synapses isolated from Alzheimer’s disease and control subject brains identifies over 5,500 proteins in human synapses.
- In silico analysis reveals region-specific decreases in proteins involved in synaptic and mitochondrial function and increases in proteins involved in neuroimmune signaling and intracellular signaling in AD.
- The apolipoprotein E4 risk gene is associated with exacerbated changes in synaptic proteins in AD.

## Introduction

Dementia poses one of the biggest societal challenges of the 21^st^ century. Over 50 million people are living with dementia worldwide, it costs over $800 billion per year to care for them, and there are currently no disease modifying treatments ^1^. One of the barriers to developing effective therapies for Alzheimer’s disease, the most common cause of dementia, lies in the lack of a comprehensive understanding of the brain changes that cause neurodegeneration. Extracellular amyloid beta (Aβ) plaques, intracellular neurofibrillary tangles composed of hyperphosphorylated tau protein, and severe brain atrophy are the major neuropathological hallmarks of AD ^2^. There are many genetic risk factors for developing sporadic AD, the strongest of which is inheritance of the apolipoprotein E epsilon 4 allele (*APOE*4). Inheritance of one copy of *APOE*4 is associated with a 3 fold increase in disease risk and inheritance of two copies with an over 10 fold increased risk ^3^. In addition to its known roles in Aβ production and clearance, we have observed that apoE protein is involved in Aβ-mediated synapse degeneration ^4, 5^, which is important as synapse loss is the strongest pathological correlate with cognitive decline in AD ^6, 7, 8^. ApoE4 causes more synaptic loss around plaques when expressed in mouse models of familial AD ^4^. Using high resolution imaging in human post-mortem brain tissue, we similarly observed exacerbated synapse loss in *APOE*4 carriers and further that apoE4 is associated with more accumulation of oligomeric Aß at synapses ^5^. More recent data implicates *APOE*4 in tau mediated neurodegeneration ^9^ and also inflammatory *TREM2* mediated microglial phenotypes ^10^, which may be important for synapse degeneration.

Recent data from postnatal human brain samples shows that proteomic datasets can reveal differences in proteins that are not observed in RNA expression data, arguing the importance of building strong resource datasets at the level of protein in human diseases ^11^. Thus far there have been several proteomic studies of human AD brain tissue (supplemental table 1), but a comprehensive dataset on human synaptic proteins examining the downstream molecular effects of *APOE* genotype in AD remains unavailable.

**Table 1:** Subject characteristics.

| MRC BBN number | group | ApoE genotype | sex | age | PMI (h) | brain weight (g) | brain pH | Braak Stage | neuropath diagnosis and co-morbidities |
| --- | --- | --- | --- | --- | --- | --- | --- | --- | --- |
| BBN_14395 | control | 3/3 | f | 74 | 41 | 1520 | 6.3 | 0 | control, Mild degree of small vessel disease |
| BBN_19686 | control | 3/3 | f | 77 | 75 | 1320 | 6.5 | I | control, mild Alzheimer's Disease pathology, sCAA, Lacunar vascular disease |
| BBN_20122 | control | 3/3 | m | 59 | 74 | 1500 | 6.1 | 0 | control, No significant abnormalities |
| BBN_22612 | control | 3/3 | m | 61 | 70 | 1300 | 6.1 | 0 | control, No significant abnormalities |
| BBN_24340 | control | 3/3 | m | 53 | 53 | 1400 | 6.5 | 0 | control, Significant atherosclerosis in larger vessels, mild small vessel disease |
| BBN001.26495 | control | 3/3 | m | 78 | 39 | 1290 | 6.17 | I | control, mild Alzheimer's Disease pathology |
| BBN001.28402 | control | 3/3 | m | 79 | 49 | 1503 | 6.33 | I | control, mild Alzheimer's pathology, mild WM pathology, Moderate non-amyloid SVD, Encephalopathy, hepatic |
| BBN001.28406 | control | 3/3 | m | 79 | 72 | 1437 | 6.13 | II | control, mild Alzheimer's pathology, Moderate arteriolar CAA, WM pathology, Mild non-amyloid SVD |
| BBN001.28793 | control | 3/3 | f | 79 | 72 | 1219 | 5.95 | II | control, mild Alzheimer's pathology, moderate WM pathology, moderate non-amyloid SVD |
| BBN_15258 | dementia | 3/3 | m | 65 | 80 | 1335 | 6.1 | VI | Alzheimer's Disease, LBD neocortical subtype |
| BBN_19595 | dementia | 3/3 | m | 87 | 58 | 1420 | 6.5 | VI | Alzheimer's Disease, CAA, SVD with lacunar infarcts |
| BBN_19994 | dementia | 3/3 | f | 87 | 89 | 1270 | 5.9 | VI | Alzheimer's Disease, sCAA, Vascular disease lacunar, thrombus, embolus |
| BBN_22223 | dementia | 3/3 | f | 87 | 83 | 1200 | 6.7 | IV | Alzheimer's Disease, cerebral haemorrhage, vascular disease lacunar |
| BBN_24527 | dementia | 3/3 | m | 81 | 74 | 1160 | 6.1 | VI | Alzheimer's Disease, Vascular Disease lacunar |
| BBN001.28410 | dementia | 3/3 | f | 62 | 109 | 1029 | 6.04 | VI | Alzheimer's Disease, Mild arteriolar CAA, Moderate non-amyloid SVD |
| BBN001.28771 | dementia | 3/3 | m | 85 | 91 | 1183 | 5.95 | VI | Alzheimer's disease, Severe arteriolar CAA, WM pathology moderate, Moderate non-amyloid SVD |
| BBN_28785 | dementia | 3/3 | f | 78 | 76 | 960 | 5.9 |  | Corticobasal degeneration FTLD, WM pathology, mild. Moderate non-amyloid SVD |
| BBN_15221 | control | 3/4 | m | 53 | 114 | 1650 | 6.1 | 0 | control, No significant abnormalities |
| BBN_15809 | control | 3/4 | m | 58 | 90 | 1470 | 5.9 | 0 | control, mild small vessel disease |
| BBN_16425 | control | 3/4 | m | 61 | 99 | 1270 | 6.2 | 0 | control, evidence of cerebrovascular disease, no infarcts |
| BBN_20593 | control | 3/4 | m | 60 | 52 | 1460 | 6 | 0 | control, no significant abnormalities |
| BBN_20120 | control | 3/4 | m | 53 | 97 | 1400 | 6.4 | 0 | control, no significant abnormalities |
| BBN_22629 | control | 3/4 | f | 59 | 53 | 1280 | 6.3 | 0 | control, no significant abnormalities |
| BBN_001.2555 | control | 3/4 | m | 74 | 66 | 1350 | 6.3 | 0 | control, small vessel lipohyalinosis, large vessel atherosclerosis |
| BBN_10591 | dementia | 3/4 | m | 86 | 76 | 1470 |  | VI | Alzheimer's Disease, small vessel disease |
| BBN_15810 | dementia | 3/4 | f | 73 | 96 | 1090 | 6.2 | VI | Alzheimer's Disease, vascular disease |
| BBN_001.15811 | dementia | 3/4 | f | 81 | 41 | 1457 | 6.3 | VI | Alzheimer's Disease, sCAA, Intracerebral haemorrhage, vascular disease |
| BBN_19690 | dementia | 3/4 | m | 57 | 58 | 1200 | 5.9 | VI | Alzheimer's disease, |
| BBN_23394 | dementia | 3/4 | f | 88 | 59 | 1165 | 6.3 | VI | Alzheimer's Disease, sCAA, Intracerebral haemorrhage, vascular disease lacunar |
| BBN_001.24322 | dementia | 3/4 | m | 80 | 101 | 1410 | 6 | VI | Alzheimer's Disease, sCAA |
| BBN_24526 | dementia | 3/4 | m | 79 | 65 | 1300 | 6.05 | VI | Alzheimer's Disease |
| BBN_25739 | dementia | 3/4 | f | 85 | 45 | 1375 | 5.77 | VI | Alzheimer's Disease, sCAA, focal TDP43 within the entorhinal cortex. |
| BBN_001.26718 | dementia | 3/4 | m | 78 | 74 | 1367 | 6.13 | VI | Alzheimer's disease, moderate non amyloid arteriolar CAA |
| BBN_26712 | dementia | 3/4 | m | 76 | 66 | 1467 | 6.48 | VI | Alzheimer's disease, Moderate non amyloid arteriolar SVD, Severe arteriolar CAA, Limbic Lewy body disease |

In order to further our understanding of how *APOE* may be influencing synaptic vulnerability in AD, we have performed a comprehensive proteomic study of human post-mortem brain tissue through a series of molecular comparisons allowing us to assess the relative contribution of both regional vulnerability and *APOE* variants to AD pathogenesis. Although our study is in postmortem tissue which has inherent limitations including looking at a snapshot of the end stage of the disease, the inclusion of a less affected brain region allows some novel insight into changes that may be occurring in synapses earlier in the degenerative process. We provide a unique proteomic resource identifying over 5,500 proteins in human synaptoneurosome preparations. Additionally, we highlight multiple proteins and molecular pathways that are modified in AD with brain region and *APOE* genotype status. *In silico* analysis reveals that proteins involved in glutamatergic synaptic signalling and synaptic plasticity are decreased in AD with temporal cortex (which has high levels of pathology) being more severely affected than occipital cortex (which has lower levels of pathology) and *APOE*4 carriers more affected than *APOE*3 carriers. Alterations in glial proteins important for neuroimmune signalling were also detected using *in silico* analysis, and further investigation revealed a host of proteins involved in the complement cascade are not only found in human synapses but are increased in AD compared to control brain. In addition to providing a resource for the field, our data support the hypothesis that *APOE* genotype plays an important role in synaptic dysfunction and degeneration in AD. The proteins and pathways identified as altered in this study can now be investigated in more detail for their potential targeted therapies to delay or prevent synaptic alterations and the consequential symptoms contributing to dementia.

## Results

### Development of a human post-mortem synaptic reference proteome

To better understand the changes in synapses that may contribute to disease pathogenesis in AD and how the genetic risk factor *APOE* contributes to synaptic vulnerability, we conducted a comprehensive proteomic study of human post-mortem brain tissue. For this study, we examined two brain regions, superior temporal gyrus (BA41/42) which has a severe pathological burden and primary visual cortex (BA17) which is less severely affected even at the end stages of disease (Figure 1A & B ^2^). With this study design incorporating disease, brain region, and APOE genotype, it is possible to design a series of comparisons which will enable the interrogation of complex proteomic comparisons in a biologically meaningful way (Figure 1C). Through the MRC Edinburgh sudden death brain bank, we were able to access samples from 34 brain tissue donors whose condition and underlying genetics were amenable to this particular investigation. Details of subjects in the study can be found in Table 1.

**Fig. 1:**
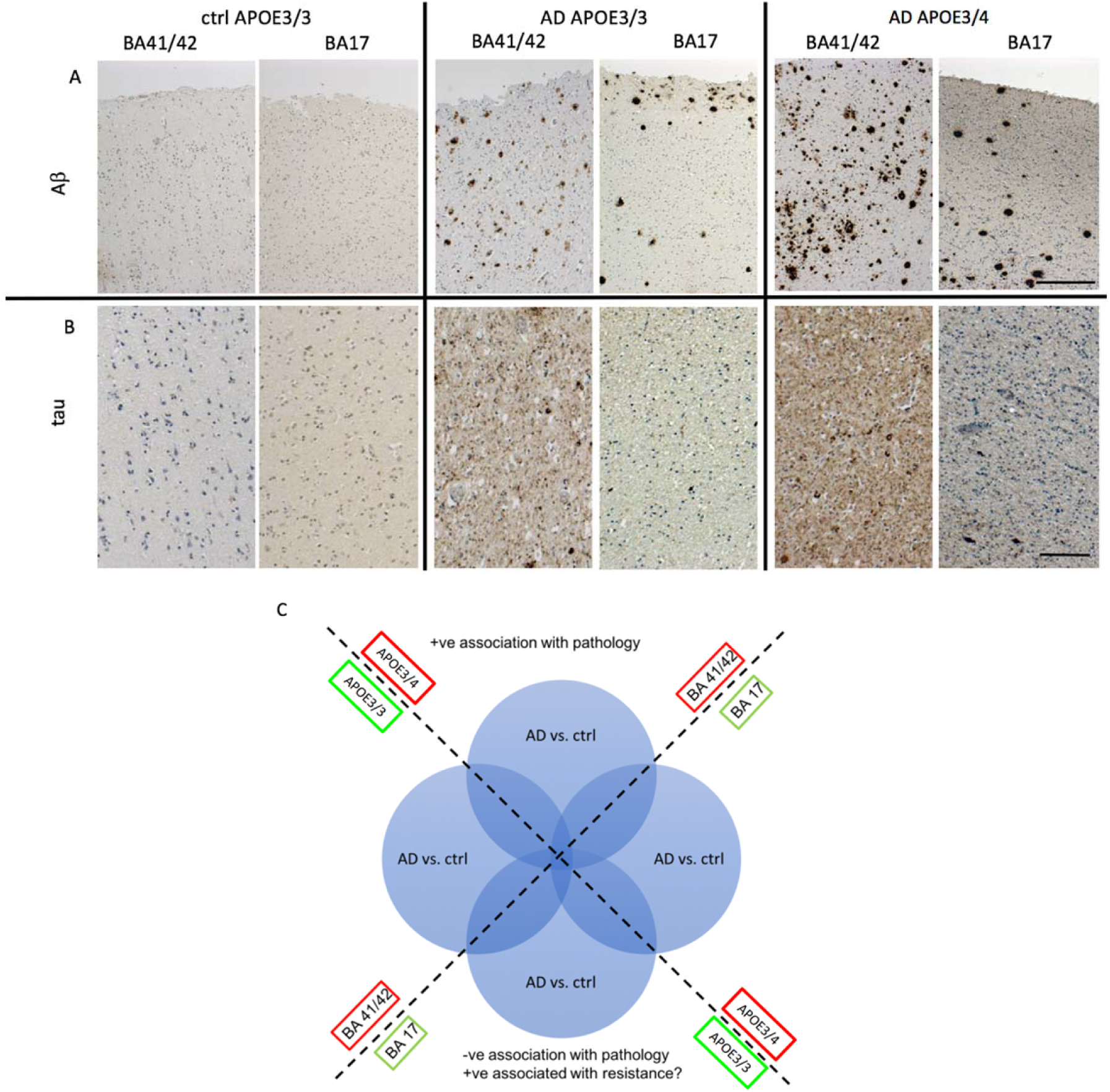
Increased pathology in temporal compared to occipital cortex and in *APOE4* compared to *APOE3* carriers. Representative images of immunohistochemistry for amyloid beta (A) and tau (B) highlight the higher pathological load in *APOE*4 carriers than *APOE*3 carriers and more pathology in superior temporal gyrus (BA41/42) compared to primary visual cortex (BA17). For analysis, we examined ratios of AD vs control samples in the 2 brain regions and with *APOE*3/3 or *APOE*3/4 genotype to examine how the synaptic proteome changes with differential pathology (C). Scale bars represent 200µm (A) 50µm (B).

Synaptic preparations were prepared and quality control for post-mortem protein degradation were confirmed as previously described^12, 13, 14^ (Supplementary Figure 1 for example total protein stains and western blotting). Any synaptoneurosome preparations containing nuclear histone protein were discarded and fresh preparations made from the same case. Protein degradation was assessed using the “HUSPIR” ratio or degradation index ^15^. To find this value, synaptoneurosmes run on western blot were probed using an antibody against NMDAR2B which recognizes two bands, the full-length protein at 170 kDa and a degradation product at 150kDa. This degradation only occurs post-mortem as it is not found in biopsy tissue thus comparing these two bands is an accepted indication of post mortem synaptic protein degradation. The degradation index was found to significantly correlate with RNA integrity number (p=0.009, R^2^=0.215, linear regression analysis) and lower but still significant correlation with brain pH (p=0.023, R^2^=0.184, linear regression analysis) but not at all with Post Mortem Interval. RIN and pH are routinely collected for all brain bank samples and used as a proxy for tissue integrity. Here we confirm that protein degradation correlates better with these markers than with post-mortem interval highlighting the importance of tissue handling in maintaining protein integrity.

Having confirmed that the extracted protein is of appropriate quality, we then applied a comprehensive workflow to enable us to assess (at the protein level) the relative contribution of both regional vulnerability and *APOE* variants as a risk factors to AD pathogenesis (Figure 1C, Figure 2) ^12, 13^. We performed a complex 8-plex TMT LC-MS/MS analysis. Representative pools based on the outlined groupings were generated from these synaptic protein extracts. By pooling individual samples according to APOE genotype and cortical area, we were able to reduce potential noise in the system generated through inter individual differences, subtle post mortem handling differences and/or sample isolation ^16^. Thus, 25 μg of each of the 34 subjects were pooled according to disease status, APOE genotype (3/3 or 3/4) and brain region (BA41/42 or BA17). The inclusion of an equivalent proportion of each protein isolate into a readily comparable pool allowed the generation of a molecular fingerprint representative of each condition and enables subsequent analysis of individual patient variability in the resulting validatory work (as a deviation from the population signal, as previously described ^14, 17^).

**Fig. 2:**
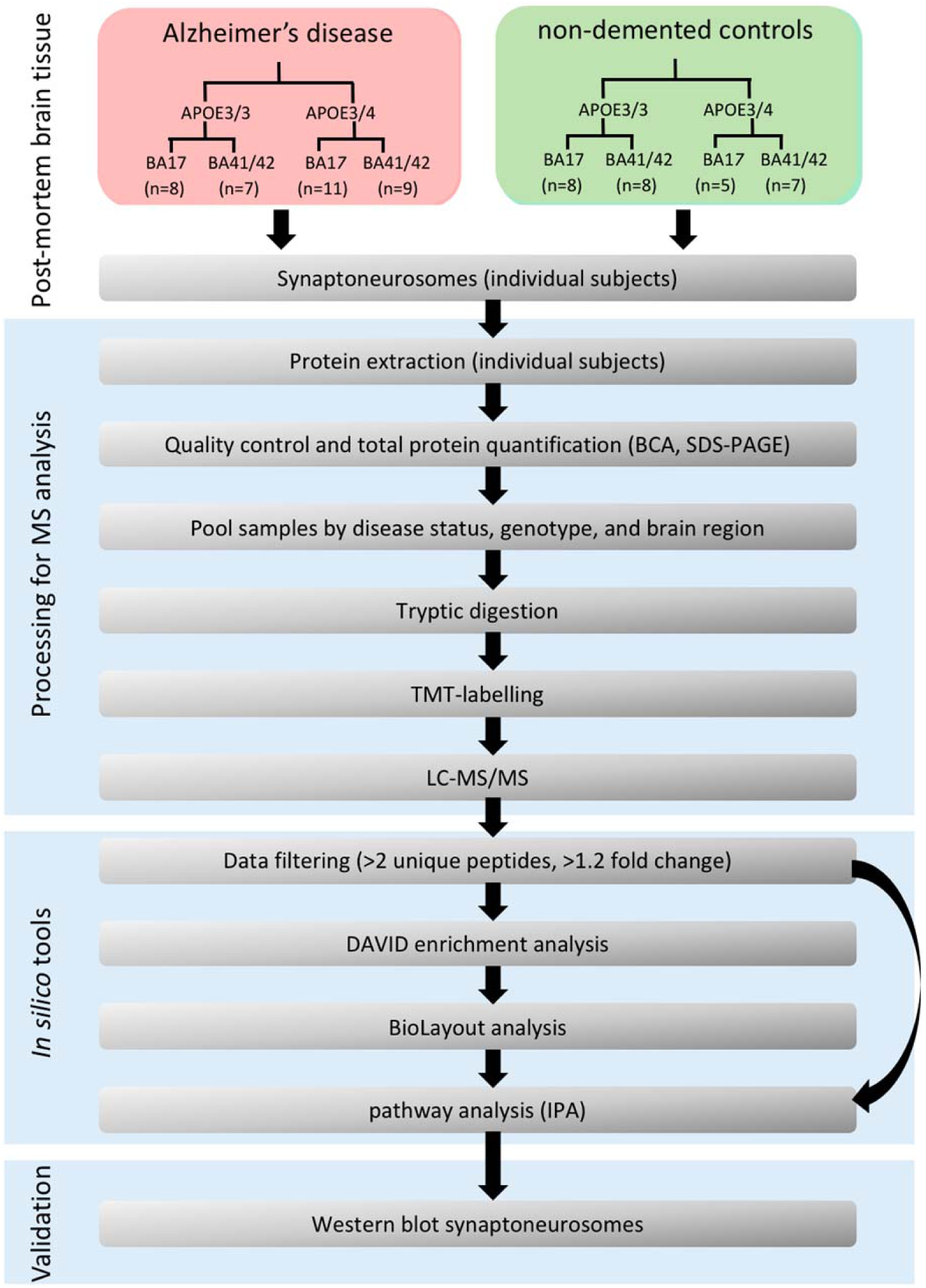
Workflow. Samples were prepared from postmortem tissue and processed for proteomics analysis according to the workflow shown.

**Supplemental Figure 1:**
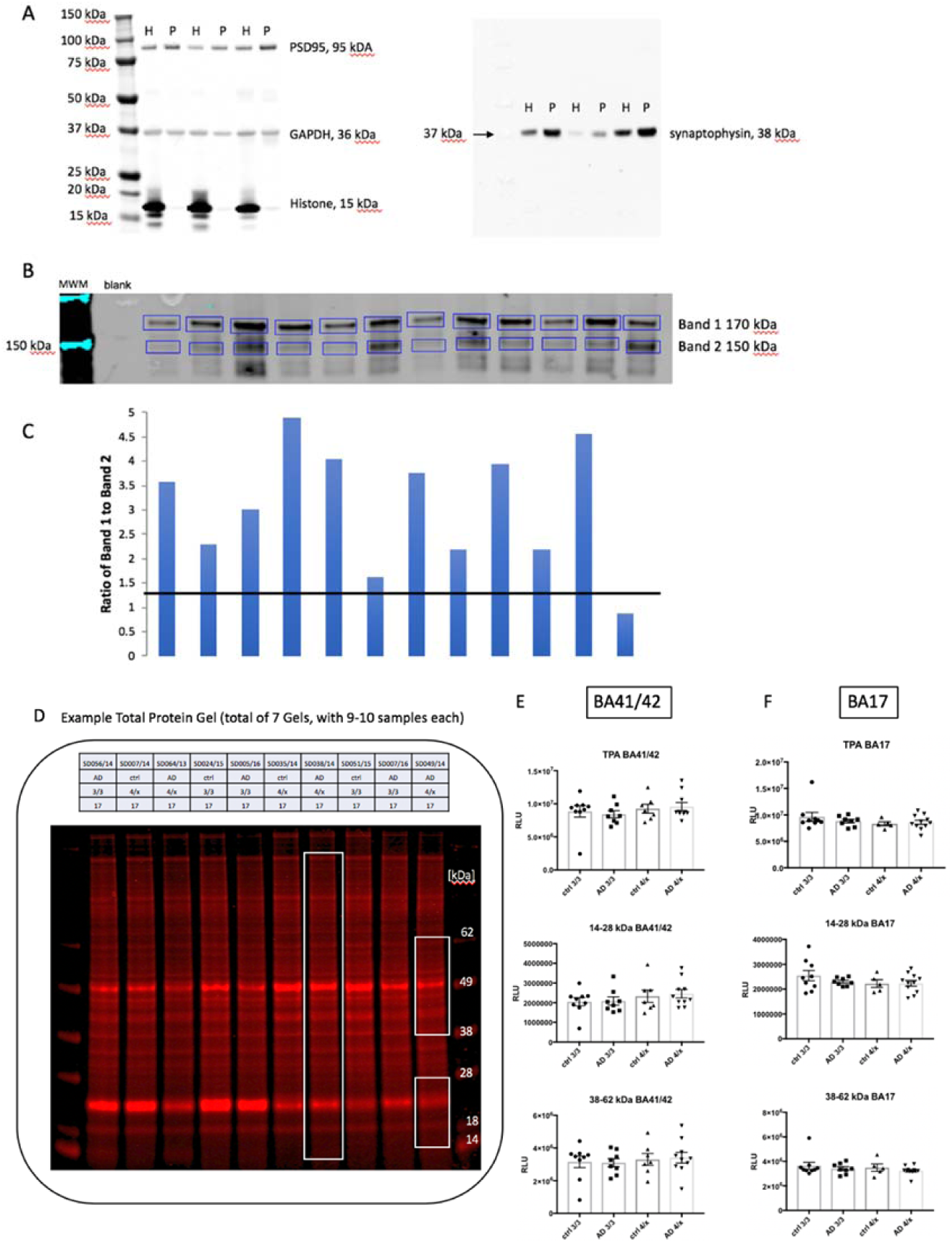
Enrichment and integrity analysis of synaptic proteins. A) A representative western blot from 3 cases shows the enrichment of synaptic proteins and exclusions of histones from the synaptoneurosome preparation (P) compared to crude homogenate (H) protein from that sample. Blots were probed for PSD95, synaptophysin, histone and GAPDH. The level of the synaptic proteins varies between the preps, however the enrichment of both synaptic proteins appears to be similar between preparations. B) Post-mortem degradation was assessed by western blotting for NMDA NR2A subunit. C) HUSPIR degradation index was calculated as the ratio of band 1 to band 2 and any cases falling below a ratio of 1 (black line) were excluded. Total protein analysis was also used to determine whether any samples showed evidence of protein degradation. Boxes in panel (D) indicate the molecular weight ranges analysed for total protein stain. Quantification reveals no difference in total protein in BA41/42 (E) or BA17 (F) samples (One way ANOVAs, p>0.05).

Following detection on the mass spec, quantitation of the dataset using MaxQuant ^18^ and Andromeda search engine software ^19^ we identified 7148 protein identifications (IDs) in total. These 7148 identified proteins were then filtered to include only proteins identified by 2 or more unique peptides (Figure 3C). This yielded 5678 proteins. A DAVID enrichment analysis of this filtered list of proteins served to further confirmed enrichment of the samples for synaptic material (Supplemental table 2).

**Fig. 3:**
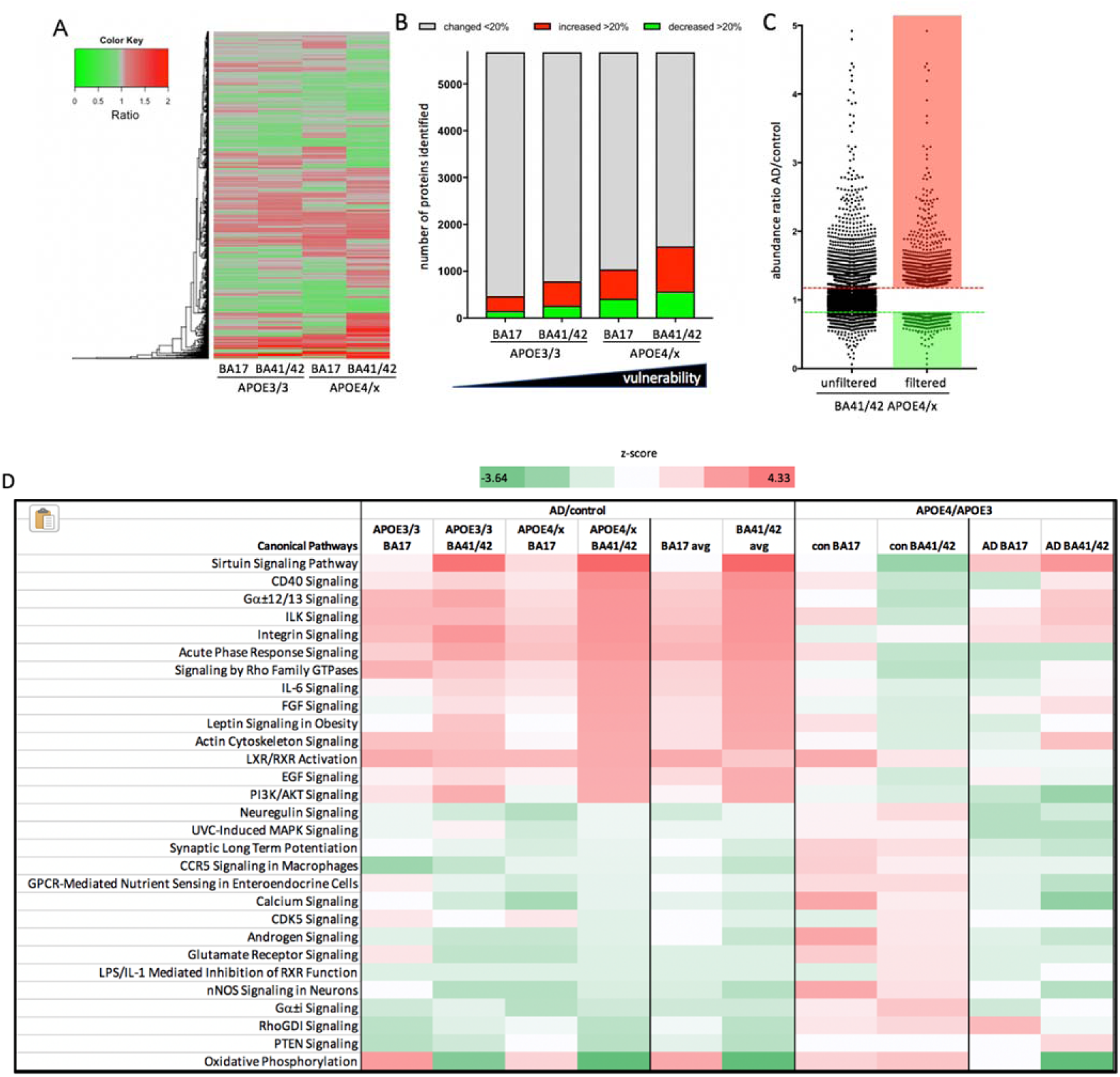
Synaptic proteomes are altered by AD and *APOE* genotype. A) Heat map with hierarchical clustering demonstrates the differential abundance of the 5678 individual proteins across “spared” and “vulnerable” brain regions and with *APOE* genotype. The ratios compare AD to control in each condition. B) Stacked bar chart demonstrates there is a direct correlation between the number of proteins that are differentially expressed, by 20% or more (up or down regulation), with the “vulnerability status’ of the synapses as determined by both genotype and brain region. C) Graphical representation of protein abundance ratios in the comparison AD vs. ctrl (*APOE*3/4, BA41/42) before (left) and after applying the filters of at least 2 unique peptide IDs and >20% change between AD and control (right). The dotted lines indicate a change of 20% up or down (note 4 proteins out of 5678 which have abundance ratios higher than 5 are excluded from the graph in panel C). D) Using Ingenuity Pathway analysis to investigate pathway alterations in the 1532 proteins that were changed more than 20% in the most vulnerable brain condition (*APOE*3/4 BA41/42) shows that in AD there is a clear upregulation of pathways involved in the immune response and cellular signaling and down regulation of several pathways involved in synaptic function including long term potentiation, glutamate signaling, and calcium ignaling. These pathways were differentially regulated in the different brain regions and *APOE* genotypes. Comparing *APOE*4 to *APOE*3 carriers within the control (con) and AD groups similarly reveals region-specific effects of *APOE* isoform on the synaptic proteome. Z-scores are plotted in D with upregulated proteins shown in red and downregulated in green.

**Supplemental table 2:** DAVID Analysis confirms enrichment of synaptic proteins.

| Term | P-Value | Fold Enrichment | Bonferroni | Benjamini |
| --- | --- | --- | --- | --- |
| clathrin coat of coated pit | 3.33E-03 | 3.4 | 9.80E-01 | 2.58E-02 |
| mitochondrial alpha-ketoglutarate dehydrogenase complex | 7.87E-02 | 3.4 | 1.00E+00 | 3.73E-01 |
| synaptobrevin 2-SNAP-25-syntaxin-1a-complexin I complex | 7.87E-02 | 3.4 | 1.00E+00 | 3.73E-01 |
| calcineurin complex | 7.87E-02 | 3.4 | 1.00E+00 | 3.73E-01 |
| mitochondrial respiratory chain complex I | 2.90E-16 | 3.0 | 3.89E-13 | 1.30E-14 |
| mitochondrial proton-transporting ATP synthase complex | 1.21E-05 | 2.8 | 1.40E-02 | 1.53E-04 |
| postsynaptic density | 2.23E-41 | 2.6 | 2.60E-38 | 2.60E-39 |
| excitatory synapse | 3.21E-05 | 2.6 | 3.68E-02 | 3.86E-04 |
| dendritic spine | 3.51E-21 | 2.6 | 4.10E-18 | 2.05E-19 |
| presynaptic active zone | 2.00E-06 | 2.6 | 2.33E-03 | 2.99E-05 |
| synaptic vesicle | 9.35E-19 | 2.6 | 1.09E-15 | 4.37E-17 |
| AMPA glutamate receptor complex | 4.98E-06 | 2.6 | 5.80E-03 | 6.84E-05 |
| synaptic vesicle membrane | 2.16E-10 | 2.5 | 2.52E-07 | 5.04E-09 |
| presynaptic membrane | 1.43E-11 | 2.5 | 1.67E-08 | 4.07E-10 |

Having confirmed that the proteomic data is likely to be representative of the synaptically enriched starting material we then filtered to include only those demonstrating differential abundance of equal to or greater twenty percent (up or down regulated) in the comparison AD vs. ctrl in BA41/42 in APOE4 carriers (Fig. 3C). After applying these further two data filtering steps we obtained a set of 1532 protein IDs identified with high confidence and meeting our differential abundance criteria (Figure 3C).

### *In-silico* analysis revealed differences in abundance ratios correlating with increasing vulnerability for AD neuropathology

To determine potential differences in the synaptic proteomes of AD patients vs. control subjects dependent on *APOE* genotype and brain region, we focused on the protein abundance ratios as calculated by dividing values from AD patients by matched control subjects, subcategorised for *APOE*3/4 or *APOE*3/3 genotype, and segregated by brain region (Fig. 3A-C). More proteins were changed in AD patients compared to controls in BA41/42 of *APOE*4 carriers than any other condition. The numbers of protein changes increases progressively from *APOE*3/3 BA17 < *APOE*3/3 BA41/42 < *APOE*3/4 BA17 < *APOE*3/4 BA41/42. Looking at the 15 most upregulated and downregulated pathways detected by Ingenuity Pathway Analysis software (Fig. 3D) reveals that several pathways involved in intracellular signalling, glial proteins involved in glia-neuron interactions, and the immune response are upregulated in AD compared to control with generally larger effects in the temporal cortex BA41/42. Sirtuin signalling is the most upregulated pathway in AD vs control with higher levels in BA41/42 and in *APOE*4 carriers. Sirtuin 2 is increased 17% and sitruin 3 by 28% and Bax and CPS1, which are involved in sirtuin signaling in mitochondria are increased 28% and 43% respectively in AD *APOE*3/4 BA/41/42 compared to controls. Downregulated pathways include many involved in synaptic function such as synaptic long term potentiation, glutamate signalling, and calcium signalling. The most downregulated pathway in AD *APOE*3/4 BA/41/42 was oxidative phosphorylation including significant downregulation of proteins in complex I, IV, and V. This pathway was increased in BA17 indicating a potential compensatory effect in BA17 which has less pathology at end stages of disease. This combination of region specific decreases in synaptic and mitochondrial proteins is very interesting in light of our recent paper showing decreased numbers of mitochondria in synaptic terminals in BA41/42 using electron microscopy ^20^. All pathways detected with IPA analysis are available in supplemental table 3.

Our previous work examining synapses in human post-mortem tissue has revealed that the handling of the body and the tissue is critical for maintaining structural and molecular integrity of synapses ^20, 21^. In particular, we observe that rapid cooling of the body after death preserves synapse structure and molecular integrity better even than short post-mortem intervals. Therefore in this study we used samples only from the Edinburgh MRC Sudden Death Brain Bank whose robust handling protocol is carried out on each individual ensuring that the data is as comparable as possible. This precluded precise age and sex matching of our control groups with *APOE*3/3 and *APOE*3/4 genotypes (see table 1), which could mean that there are confounding effects of age and sex on our AD/control comparisons. However, when we compare BA41/42 of the AD *APOE*3/4 to AD *APOE*3/3 cases, which are reasonably age and sex matched, we observe many of the same pathways increased in AD *APOE*4 carriers compared to AD *APOE*3 carriers as those we observed in the comparison of AD vs control *APOE*4 carriers (Fig. 4D). Less changes were observed in BA17 from AD *APOE*4 carriers compared to AD *APOE*3 carriers. This strongly supports our conclusion that *APOE*4 influences the synaptic proteome in AD in a region-specific manner. Interestingly, in non-demented controls when *APOE*4 carriers are compared to *APOE*3 carriers in BA41/42, there are changes in the opposite direction to those that are observed in E4 AD vs control.

**Fig. 4:**
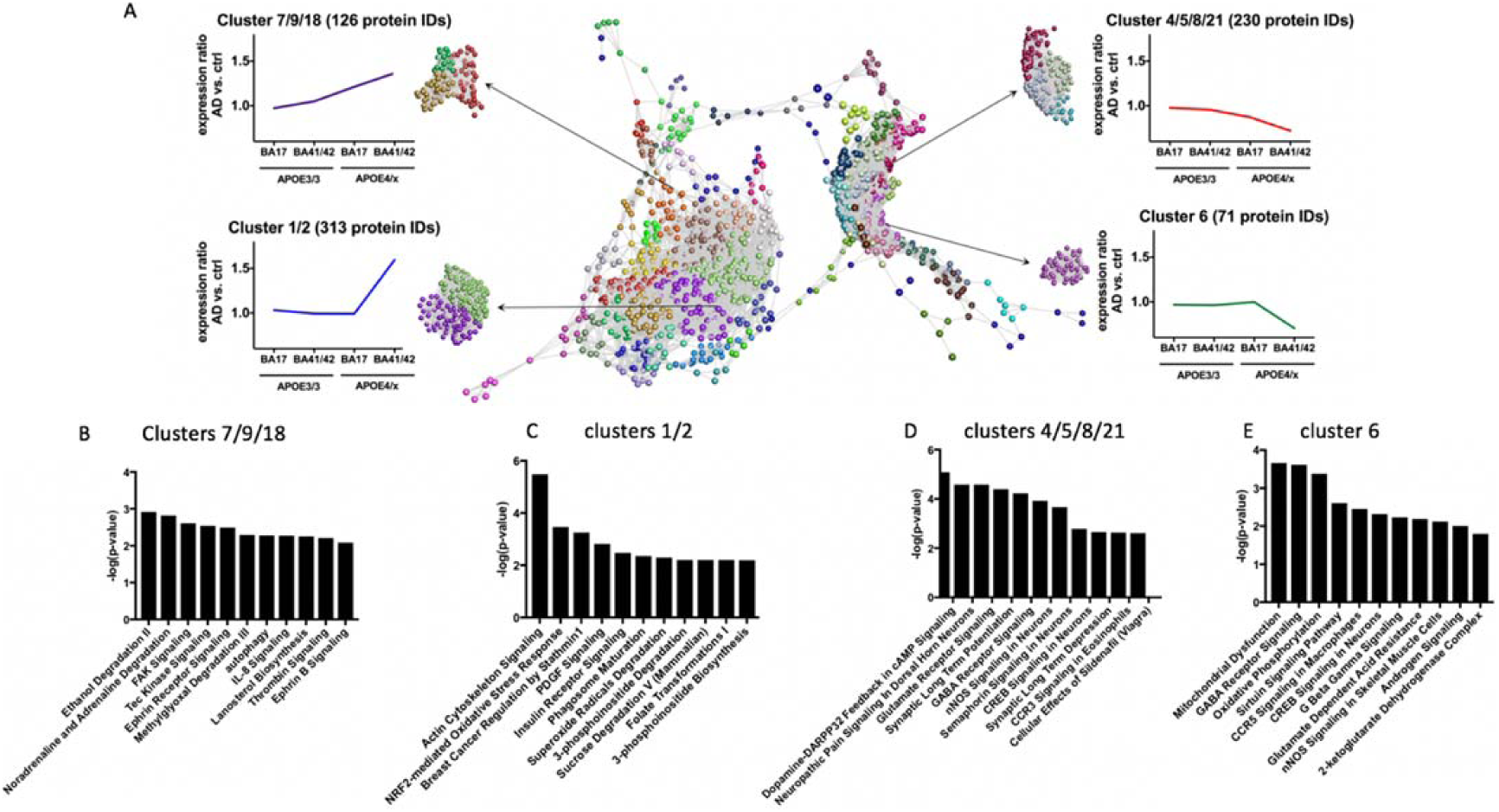
Clusters of protein changes. (A) Graphia Professional representation of proteomic abundance data across differentially vulnerable synaptic populations. Each sphere represents a single protein and the edge represents how similar their abundance trend is towards the other proteins in the dataset. The closer the spheres are the more similar the abundance trend. The colours represent the different clusters formed by grouping proteins with similar abundance trends. The resulting profiles were grouped into four different categories as shown in the example graphical abundance trends for further analysis. The annotation associated with each graph represents the cluster numbers which fit each trend and the number of associated proteins. Graphs B-E show the top pathway changes in the clusters indicated.

To further examine trends in protein changes in an unbiased manner, we performed clustering analysis to detect differences in the abundance ratios across these differentially vulnerable synaptic populations. Proteins were clustered according to their abundance profile across the four calculated ratios (Fig. 4). Each sphere represents an individual protein ID and the distances between the spheres indicate the similarity in abundance profile. Different colours are used to group proteins together in clusters based on abundance profile similarity. In order to analyse the impact/influence of *APOE* genotype and cortical region, we focused specifically on clusters showing a steady increase or decrease in protein abundance across the four groups or only demonstrating differences in protein abundance in *APOE*- ε3/4 carriers in BA41/42 (Fig. 4). In order to determine if these abundance profile specific clusters were associated with specific pathways and/or canonical cascades, we have carried out a higher order functional clustering analysis using IPA software. IPA pathway analysis and DAVID enrichment analysis highlighted multiple affected pathways. In clusters 4, 5, 8 and 21 which have progressive decreases in AD/control ratios as pathological severity increases (Fig 4), we observe pathways involved in synaptic function are decreased including glutamate signalling, synaptic long term potentiation, GABA receptor signalling, CREB signalling, and synaptic long-term depression. Cluster 6 containing proteins that were sharply decreased in the condition with most pathology, AD *APOE*4 BA41/42, similarly showed decreases in pathways involved in synaptic function including CREB signalling and GABA receptor signalling, along with decreases in pathways implicated in mitochondrial function including oxidative phosphorylation (Fig 4).

Interestingly, when examining clusters 7, 9, and 18 which were progressively increased with pathological vulnerability, we observe proteins involved in autophagy and chemokine signalling were progressively upregulated in conditions of higher synaptic vulnerability (Fig 4,). Clusters 1 and 2, which included proteins highly upregulated in AD *APOE*4 BA41/42 compared to the other groups indicate increases in pathways involved in actin cytoskeleton signalling, NRF2 mediated oxidative stress response, PDGF signalling, and insulin receptor signalling, which all implicate non-neuronal contributors to synapse degeneration as has been recently emphasised for AD risk by genetic studies ^22^.

Along with the unbiased bioinformatic analyses, we further interrogated our proteomics dataset to examine proteins of interest based on what is known about synapse degeneration in AD from model systems. In addition to loss of proteins involved in synaptic function in remaining synapses in AD as shown with bioinformatics, it is likely that there is some degree of synaptic remodelling/compensation taking place as some synaptic proteins were increased. The synaptic receptor TMEM97 is increased 2.8 fold in remaining synapses in AD vs control *APOE*4 carriers in BA41/42 (Fig 5). This is particularly interesting because TMEM97 is the sigma 2 receptor, and compounds that disrupt interaction of Aβ and sigma 2 receptors are protective in mouse models and are being tested for efficacy in human AD as a therapeutic ^23^.

**Figure 5:**
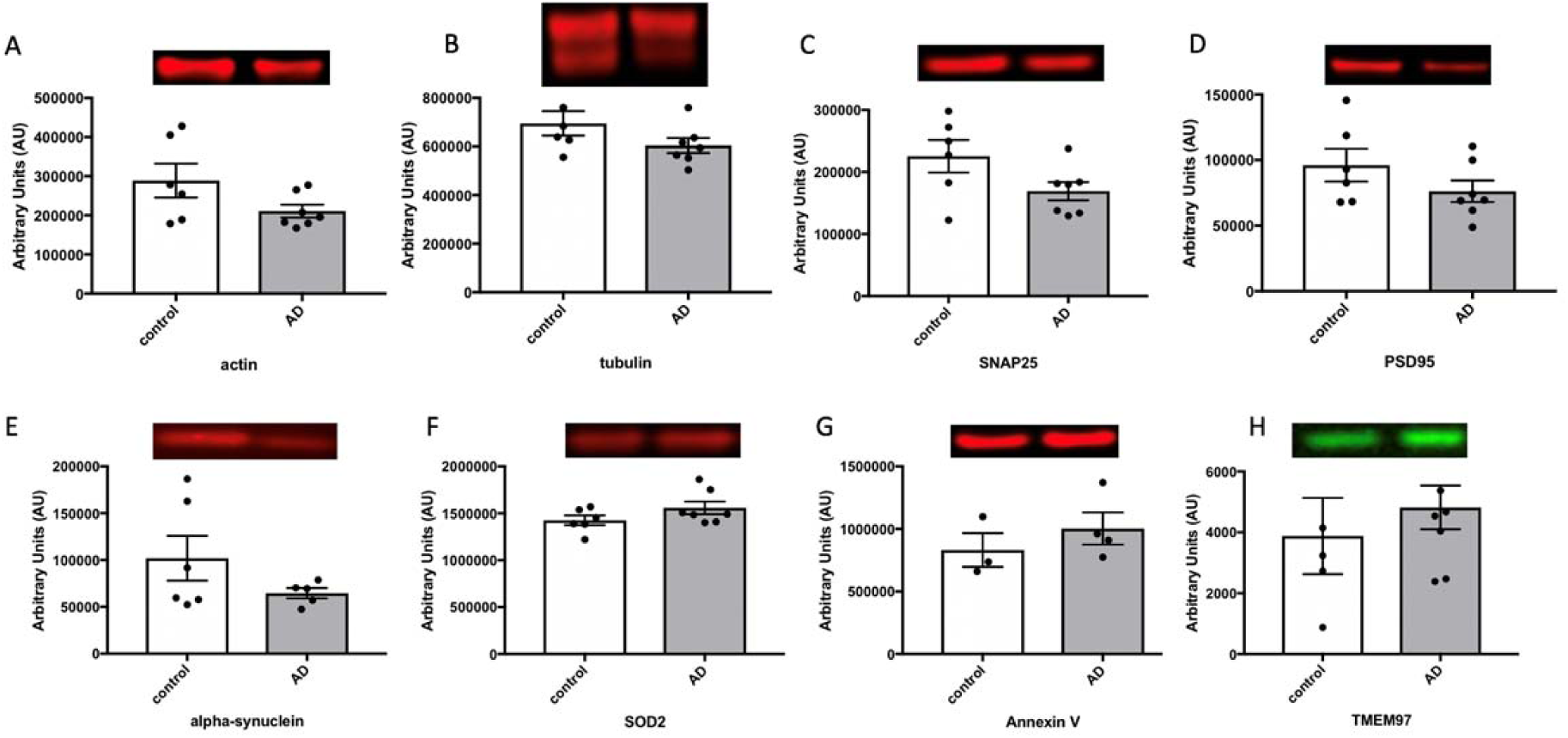
Western blot validation of proteomic analysis. Quantitative fluorescent western blotting was used in BA41/42 of *APOE*4 cases to confirm changes in proteins seen with proteomics. Western blot validation from non-demented control (white bars) and AD cases (grey bars) show examples of proteins that were decreased unchanged, and increased in AD. Data are shown as mean and error bars represent the standard error of the mean. Each dot represents the value from a single case. Housekeeping proteins actin (**A**) and tubulin (**B**), commonly used loading controls, are downregulated in AD patients. **C, D.** Pre synaptic proteins (SNAP25) and post-synaptic (PSD95) respectively are downregulated in the synapses of AD patients. **E.** Abundance of alpha synuclein, a presynaptic protein that is often associated with neurodegeneration, is also appreciably decreased in AD patients. **F.** Mitochondrial protein SOD2 is unchanged in AD vs control. **G.** Annexin V is upregulated in the synapses of AD patients. **H.** TMEM97, a proposed synaptic receptor for Abeta is increased in AD. Total protein stained gels were used as a loading control as described in ^29^. Data are shown as means and standard errors with each dot representing an individual case. Non-cropped blots are found in supplementary figure 3.

Recent data strongly implicate the complement cascade and microglia in Aβ mediated synapse loss in mouse models of amyloid deposition ^24, 25^. Two studies recently demonstrated upregulation of components of the complement system in AD brain and influence of complement cascade in synapse dysfunction and loss in a mouse model of tauopathy ^26, 27^. Based on these data, we interrogated our human synaptoneurosome dataset to look at proteins important for microglial synapse phagocytosis and specifically the complement system. Pathway analysis reveals increases in CD40, IL-6, IL-8, IL-1, IL-2, IL-7 and acute phase response signalling, indicating neuroimmune signalling between neurons and glia in synapses, which is modulated by *APOE* genotype (Fig 3,4, supplemental figure 3). Further interrogation of the proteomics dataset shows increases in C1qA, B, and C in AD brain, which are most pronounced in BA41/42 of *APOE*3 carriers (over 2 fold increases). We also detect complement components C1 and C4 which are increased in most conditions AD brain without a clear effect of *APOE* genotype (supplemental table 4).

**Supplemental table 4:** Evidence for alterations in proteins involved in the complement cascade in human AD synapses.

|  | Log2<br>Normalised<br>Ratio AD/Ctrl<br>3/3 41/42 | Log2<br>Normalised<br>Ratio AD/Ctrl<br>3/4 41/42 | Log2<br>Normalised<br>Ratio AD/Ctrl<br>3/3 17 | Log<br>Normalised<br>Ratio AD/Ctrl<br>3/4 17 |
| --- | --- | --- | --- | --- |
| Complement C1q subcomponent subunit A | 1.77959139 | 0.69430641 | 1.0980274 | 0.93909556 |
| Complement C1q subcomponent subunit B | 1.44369348 | 0.68392297 | 0.7091574 | 0.91265319 |
| Complement C1q subcomponent subunit C | 1.0778657 | 0.60047611 | 0.46366574 | 0.73646894 |
| Complement C4-A | 0.84763652 | 0.73464438 | 0.89934544 | 0.66442196 |
| Complement C4-B | 0.81062454 | 0.98633814 | -0.0877303 | 0.38715829 |
| Complement C1s | 0.68375909 | 0.2130724 | 0.39700963 | 0.56568522 |
| Complement component 1 | 0.29882572 | 0.59348713 | -0.5659684 | 0.40746573 |

In order to validate the proteomics data, we selected a subset of proteins for western blot analysis in BA41/42 whose levels should be increased or decreased just past our magnitude of change cutoff as this will be more indicative of the sensitivity of the MS than highly up or downregulated proteins. We also selected a protein whose levels was unchanged to use as an internal control (Fig 5, full blots shown in supplementary figure 3).^28^

**Supplementary figure 3:**
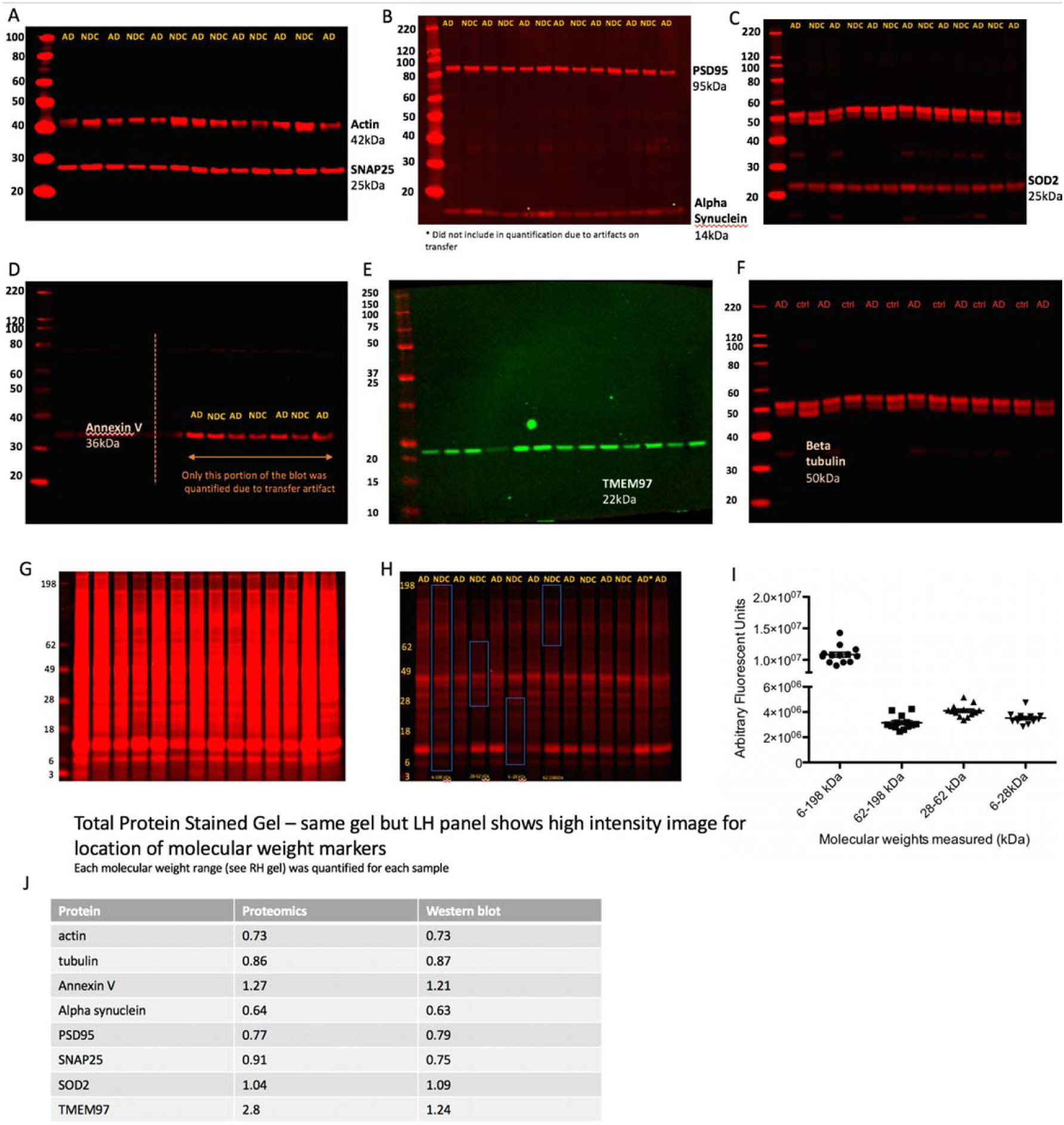
Validation western blots (uncropped) of AD vs non demented control (NDC) from BA41/42 of people with *APOE3/4* genotype. Full blots are shown for actin and SNAP25 (A), PSD95 and alpha-synuclein (B), SOD2 (C), annexin V (D), TMEM97 (E), beta tubulin (F), and total protein at high intensity (G) and low intensity (H). Each of the molecular weight ranges in H were quantified for each lane, shown in (I). Comparisons between proteomics and western blot data are shown in J.

## Discussion

Of the pathological changes associated with dementia, the best correlate to the extent of memory decline in life is the loss of synapses. Synapses are exquisitely complicated structures requiring thousands of proteins for the complex process of establishing, maintaining, and undergoing synaptic transmission. Work from animal models and human post mortem tissue indicates that synapse degeneration is a driving force in disease progression in AD. However, to date there has been a lack of data on the precise molecular changes in synapses in human AD brain, which impedes the design of hypothesis driven experiments to understand mechanisms of synapse degeneration in animal models that are likely to be relevant to human disease. In the literature, we found 19 publications using proteomics on human AD brain tissue (supplemental table 1). In proteomics studies examining whole tissue homogenates from AD brains without considering *APOE* genotype, there have been quite varied results. Three studies show decreases in synaptic proteins or pathways involved in synaptic function, which could have been explained by synapse loss ^30, 31, 32^. To our knowledge only one previous study examined the effects of *APOE* on synaptic proteins ^33^. This study used whole tissue homogenates from AD and control subjects for proteomics but focused their analysis on a group of 191 proteins that had previously been detected in synaptosome fractions of healthy subjects. With this method, they observed a downregulation of glutamate signalling proteins and an effect of *APOE*4 genotype on the abundance of these synaptic proteins. Our results significantly expand upon these findings as we biochemically isolated synaptoneurosomes from AD and control subjects and detected over 5,500 proteins, which is over 25 times more proteins examined than in the previous study of synaptic proteins. Synaptoneurosome preparations have not previously been used in proteomic studies of AD. Synaptoneurosomes, unlike other synaptic fractions, contain both the presynaptic and postsynaptic compartments, which is important due to data from model systems implicating both pre and post synapses in degenerative mechanisms. Further, the preparation might retain parts of glial processes closely associated with the synapse, which is key for understanding the role of non-neuronal cells in synapse degeneration, an important topic in the field ^22^.

We observed that multiple pathways including those associated with synaptic and mitochondrial function were downregulated with increasing vulnerability and other pathways including intracellular signalling and neuroimmune signalling proteins were increased with increasing vulnerability. This is encouraging as dysfunctional synaptic signalling is consistent with multiple lines of evidence from AD animal models^8, 34, 35^. Some of the proteins that we observe are reduced in AD synapses have been observed as CSF biomarkers associating with disease. For example, we observe a 25% decrease in neurexin 2 and over 30% decrease in neurogranin abundance in AD *APOE*4 superior temporal gyrus compared to controls, whereas recently published biomarker studies observed increases in neurexin 2 and neurogranin in CSF of people with mild cognitive impairment or AD ^36, 37, 38^. In another study of CSF, a group of 9 synaptic proteins (GluR2, GluR4, Neuroligin-2, Neurexin-3A, Neurexin-2A, Calsynytenin-1, Syntaxin-1B, Thy-1, and VAMP-2) were increased before markers of neurodegeneration are observed ^39^. Not only does this CSF data serve to further highlight the veracity of the proteomic data generated here, but critically, it also supports the notion that proteins may be removed from synapses and at least some of these cleared from the brain via CSF.

In addition to the synaptic signalling pathway changes, we observe interesting changes in immune-related signals in AD synapses. In recent work examining both iPSCs and human brain tissue, *APOE*4 was strongly associated with reduced expression of regulators of synaptic function and increased expression of microglial genes associated with the immune response ^40^. *APOE* has been shown to interact with and modulate the immune and inflammatory system in the brain especially through its interaction with another important AD genetic risk factor TREM2 ^41^. Our study also found that proteins involved in the immune system and neuroimmune signalling are dysregulated in the AD synapse. Recent evidence has indicated that the complement system of innate immunity, particularly complement components C1q and C3, are involved in synaptic death in mouse models both downstream of Aβ and tau pathology, and that ApoE forms a complex with activated C1q which blocks initiation of the complement cascade ^24, 25, 26, 27, 42^. Our study highlights the importance of these pathways in human AD brain and shows that other proteins in this cascade including complement component C4, HLA-1, and Clusterin are all increased at the synapse in AD presenting innate immunity as an attractive area for further study and therapeutic intervention. Our data indicate that within remaining synapses in human AD brain, *APOE*4 is associated with altered levels of proteins involved in synaptic function and proteins involved in the innate immune system. One possible interpretation of these data supports the hypothesis that microglia are involved in synaptic pruning during disease.

Synapses have high energy demands requiring local mitochondrial ATP production. Both Aβ and tau have been observed in model systems to impact mitochondrial function and the intracellular transport of mitochondria, which impair synaptic function ^43, 44, 45^. And there is some evidence that *APOE*4 can impair mitochondria function in cell culture.^46^ However, the role of *APOE*4 in synaptic mitochondrial function in human brain has not been studied. We observe region and *APOE* specific changes in proteins involved oxidative phosphorylation with a strong decrease in BA41/42 of AD patients that was larger in *APOE*4 carriers than *APOE*3 carriers. Conversely, this pathway was increased in BA17 with the strongest increase in E3 carriers. This could reflect a compensatory increase in mitochondrial function in brain regions that are very early in the disease process. Consistent with the BA41/42 data, we recently observed a decrease in presynaptic terminals containing multiple mitochondria in the temporal cortex of AD patients ^20^.

Synaptic degeneration is an important part of AD pathogenesis. Further understanding this process and how to delay and/or halt it may lead us towards important novel therapeutic targets not only for AD but also for other diseases for which synapse loss is an integral or important process. By coupling subcellular fractionation with anatomical knowledge of regional vulnerability and human patient genetics we were able to generate the most comprehensive synaptic proteomics profiling datasets from human AD patient samples to date. Importantly, we have determined its accuracy through experimental validation and links to existing published literature on mechanisms and biomarker identification. As an example of the potential utility of such data we were able to begin to uncover how AD and *APOE*4 impact synaptic composition and may be leading to synaptic degeneration. Although in this discussion we have highlighted specific proteins and cascades correlating with regional vulnerability and discussed their potential roles in disease progression and /or regulation this was done to highlight the potential utility of our dataset. There are many other potentially important proteins and pathways in these data to explore in future stidies. We have made the data freely available and hope that this will provide a useful resource for other researchers in the field to use at their discretion. The results described here demonstrate that *APOE* genotype has a profound impact on the molecular fingerprint of the synapse and that further understanding of the effects of these protein changes may contribute to our understanding of, and ultimately the development of novel therapies for AD.

## Methods

### Subjects

Use of human tissue for post-mortem studies has been reviewed and approved by the Edinburgh Brain Bank ethics committee and the ACCORD medical research ethics committee, AMREC (ACCORD is the Academic and Clinical Central Office for Research and Development, a joint office of the University of Edinburgh and NHS Lothian, approval number 15-HV-016). The Edinburgh Brain Bank is a Medical Research Council funded facility with research ethics committee (REC) approval (11/ES/0022).

Fresh frozen brain tissue for proteomics and paraffin embedded tissue for examination of pathology was provided from superior temporal gyrus (BA41/42) and primary visual cortex (BA17). Tissue was requested from clinically diagnosed AD and control subjects. All cases were examined by a neuropathologist, and after the proteomics results were returned, it was noted that one of the AD cases was neuropathologically classified as frotntotemporal dementia with tau-associated corticobasal degeneration (table 1).

### *APOE* genotyping

DNA was extracted from ∼25mg of cerebellum for each case using the QIAamp DNA mini kit (Qiagen, Hilden, Germany), which was used as per the manufacturer’s instructions. Polymerase chain reaction (PCR) was performed on the extracted DNA. 10μl of 2x Master mix (Promega, Madison, WI) was combined with 1μl of primer stock (20μM forward primer, 20μM reverse primer), 2μl of DMSO (Sigma-Aldrich, St Louis, MO), 6μl ddH_2_O and 1μl of isolated DNA. The forward primer was 5’taagcttggcacggctgtccaagg3’ and the reverse primer 5’acagaattcgccccggcctggtacactgcc3’ (Figure 2.2A). *APOE* ε2, *APOE* ε3, and *APOE* ε4 plasmids (generously donated by Dr E Hudry) were also amplified by PCR to use at as reference and were treated in the same way as unknown samples throughout. PCR product was digested using the restriction endonuclease HhaI (New England Biolabs, Ipswich, MA). For this 0.5μl of enzyme, 2.5μl of 10x CutSmart buffer (New England Biolabs, Ipswich, MA) and 2μl of ddH_2_O were added to each PCR reaction tube to give a total volume of 25μl. The final volume contains 50 mM Potassium Acetate, 20 mM Tris-acetate, 10 mM Magnesium Acetate, and 100 μg/ml BSA as a result of the CutSmart buffer and 10 units of HhaI. After digestion incubation 5μl of 6x Blue Loading dye (Promega, Madison, WI) containing 0.4% orange G, 0.03% bromophenol blue, 0.03% xylene cyanol FF, 15% Ficoll® 400, 10mM Tris-HCl (pH 7.5) and 50mM EDTA (pH 8.0) was added to the the reaction tube. 14μl of this mixture was then loaded onto precast 15 well Novex TBE 20% gel (Thermo Fisher Scientific, Waltham, MA) using a 25µL Hamilton Syringe. The DNA were then separated by size using electrophoresis for 2 hours at 200V. The gels were run in an XCell SureLock™ Mini-Cell (Invitrogen, Carlsbad, CA) using Novex TBE running buffer (Thermo Fisher Scientific, Waltham, MA). Bands were stained with either 2μg/ml ethidium bromide (Sigma-Aldrich, St Louis, MO) or SYBR safe DNA Gel Stain (Thermo Fisher Scientific, Waltham, MA) and visualized with an ultraviolet gel imaging system (Syngene, Cambridge, UK).

### Amyloid beta and Tau staining in human cortical sections

Tissue sections were stained for amyloid beta and tau according to a previous study ^47^. Briefly, fresh post-mortem tissue blocks were fixed in formalin and dehydrated in an ascending alcohol series. Three paraffin waxing stages were performed and 4μm thick tissue sections were cut on a Leica microtome and collected on glass slides. Immunohistochemistry for amyloid beta (BA4, M087201-2, Dako) and pTau (AT8, MN1020, Thermo) was performed using the Novolink Polymer detection system and visualized using DAB as chromogen. Images were acquired using an upright Zeiss axioImager equipped with MicroBrightfield stereology software.

### Synaptoneurosome preparation

Brain homogenates and synaptic fractions were prepared as described in Tai et al. 2012 ^48^. In brief, ∼300mg of cortical tissue was homogenized on ice in homogenization buffer (25 mM HEPES pH 7.5, 120 mM NaCl, 5 mM KCl, 1 mM MgCl_2_, 2 mM CaCl_2_, protease inhibitors (roche complete mini), phosphatase inhibitors (Millipore, 524629)). The homogenate was filtered through 2 layers of 80μm nylon filter (Millipore, NY8002500) and saved as crude homogenate. The crude homogenate was further filtered through a 5 μm filter (Millipore, SLSV025NB) and centrifuged at 1000*g* for 5 min. The pellet was washed once, then the supernatant was removed and the pellet was resuspended in in label-free buffer [100 mM Tris-HCl (pH7.6) 4% (w/v) SDS] containing 1% protease cocktail inhibitor (Thermo Fisher, UK). Homogenates were centrifuged at 20,000 x g for 20 minutes at 4°C with the soluble fraction of each sample transferred to Lo-Bind tubes (Sigma Aldrich). Protein determination using the Bicinchoninic acid assay (Pierce, UK) was carried out according to manufacturer’s guidelines.

### SDS-PAGE and western blotting

SDS-PAGE and western blotting were performed as described previously ^49^. Briefly, 5μg of protein from synaptoneurosome fractions and molecular weight marker (Li-Cor, Cambridge, UK) was loaded onto NuPAGE 4-12% Bis-Tris precast polyacrylamide 15 well gels (Invitrogen, Paisley, UK). Proteins were transferred to polyvinylidene fluoride (PDVF) membranes and blocked using Odyssey Blocking buffer (927-40,000, Li-Cor) diluted 1:1 in PBS. Primary antibodies were incubated overnight in blocking buffer and proteins were detected on an Odyssey system using 680 and 800 IR dye secondary antibodies diluted 1:10000 in blocking buffer (table 2 shows antibodies used in western blots). Total protein stains were performed with Instant Blue total protein stain per manufacturer instructions (Expedeon).

### LC-MS/MS analysis

Pools containing equal amounts of protein (25 μg per case) were prepared of each of the 8 groups (control *APOE3/3* BA41/42, control *APOE3/3* BA17, control *APOEx/4* BA41/42, control *APOEx/4* BA17, AD *APOE3/3* BA41/42, AD *APOE3/3* BA17, AD *APOEx/4* BA41/42, AD *APOEx/4* BA17). Preparation of the samples, quantification, and bioinformatics was carried out according to standardized protocols ^12, 13, 14^.

Samples were lysed in 4%SDS + 100mM tris prior to protein estimation by microBCA. Each sample was then reduced with 100 mM DTT and samples then processed using the FASP protocol ^50^ with some modifications. Samples were initially diluted 1:10 into 8M Urea and buffer exchanged to remove the SDS and tris buffer, filters were then washed 3 times with 100 mM Tris-HCL pH 8 then another 3 times with 100 mM triethyl ammonium bicarbonate (TEAB). Proteins on the filters are then digested twice at 30oc with trypsin (2 × 1ug), first overnight and then for another 6h in a final volume of 200 µl prior to addition of 200ul of 500mM NaCl. Samples were then desalted using a SPE cartridge (Empore-C18, Agilent Technologies, 7mm/3ml) and the peptides dried in a speedvac (Savant).

Desalted tryptic peptides (25µg each sample) were then dissolved in 100 µl 100 mM TEAB. The different 8 TMT labels were dissolved in 41µL of anhydrous acetonitrile, and each label then added to a different sample. The mixtures were incubated for 1 hour at room temperature and the labelling reaction was then quenched by adding 8µL of 5% hydroxylamine. Following labelling with TMT, samples were mixed, desalted using a SPE cartridge (Empore-C18, Agilent Technologies, 7mm/3ml) and the peptides dried in a speedvac (Savant). Samples were then dissolved in 200 μL ammonium formate (10 mM, pH 10) and peptides fractionated using High pH RP HPLC. A C18 Column from Waters (XBridge peptide BEH, 130Å, 3.5 µm 2.1 × 150 mm, Ireland) with a guard column (XBridge, C18, 3.5 µm, 2.1×10mm, Waters) are used on a Ultimate 3000 HPLC (Thermo-Scientific). Buffers A and B used for fractionation consists, respectively of 10 mM ammonium formate in milliQ water and 10 mM ammonium formate with 90% acetonitrile, both buffers were adjusted to pH 10 with ammonia. Fractions were collected using a WPS-3000FC auto-sampler (Thermo-Scientific) at 1 min intervals. Column and guard column were equilibrated with 2% buffer B for 20 min at a constant flow rate of 0.2 ml/min. Samples (175 µl) were loaded onto the column at 0.2 ml/min. Peptides were eluted from the column with a gradient of 2% buffer B to 5%B in 6 min then from 5% B to 60% B in 50 min. The column is washed for 16 min at 100% buffer B and equilibrated at 2% buffer B for 20 min as mentioned above. The fraction collection started 1 min after injection and stopped after 80 min (total of 80 fractions, 200 µl each). The total number of fractions concatenated was set to 20 by non-contiguous pooling and the content of the fractions dried and resuspended in 50 µl of 1% formic acid prior to analysis by nLC-MS/MS.

Analysis of peptides was performed using a Q-Exactive-HF (Thermo Scientific) mass spectrometer coupled with a UltiMate 3000 RSLCnano (Thermo Scientific) UHPLC system. nLC buffers were as follows: buffer A (2% acetonitrile and 0.1% formic acid in Milli-Q water (v/v)) and buffer B (80% acetonitrile and 0.08% formic acid in Milli-Q water (v/v). Aliquots of 15 μL of each sample (50ul in total) were loaded at 5 μL/min onto a trap column (100 μm × 2 cm, PepMap nanoViper C18 column, 5 μm, 100 Å, Thermo Scientific) equilibrated in 98% buffer A. The trap column was washed for 6 min at the same flow rate and then the trap column was switched in-line with a Thermo Scientific, resolving C18 column (75 μm × 50 cm, PepMap RSLC C18 column, 2 μm, 100 Å). The peptides were eluted from the column at a constant flow rate of 300 nl/min with a linear gradient from 95% buffer A to 40% buffer B in 122 min, and then to 98% buffer B by 132 min. The column was then washed with 95% buffer B for 15 min and re-equilibrated in 98% buffer A for 32 min. Q-Exactive HF was used in data dependent mode. A scan cycle comprised MS1 scan (m/z range from 335-1800, with a maximum ion injection time of 50 ms, a resolution of 120 000 and automatic gain control (AGC) value of 3×106) followed by 15 sequential dependant MS2 scans (with an isolation window set to 0.7 Da, resolution at 60000, maximum ion injection time at 200 ms and AGC 1×105. To ensure mass accuracy, the mass spectrometer was calibrated on the first day that the runs are performed.

The raw mass spectrometric data files obtained for each experiment were collated into a single quantitated dataset using MaxQuant 1.6.0.16 ^18^ and Andromeda search engine software ^19^ with enzyme specificity set to trypsin. Other parameters used were: (i) variable modifications, deamidation (NQ), oxidation (M), protein N-acetylation, gln-pyro-glu; (ii) fixed modifications, carbamidomethylation (C); (iii) database: uniprot-human_Sept2017 database; (iv) Reporter ion MS2 – TMT labels: TMT8plex_Nter and TMT 8plex-Lys; (v) MS/MS tolerance: FTMS-10ppm, ITMS- 0.02 Da; (vi) maximum peptide length, 6; (vii) maximum missed cleavages, 2; (viii) maximum of labelled amino acids, 3; and (ix) false discovery rate, 1%. Peptide ratios were calculated using ‘Reporter Intensity’ Data that was normalised using 1/median ratio value for each identified protein group per labelled sample.

### *In silico* analyses

Filtered data was utilised for all bioinformatics statistical analyses and filtered by the following criteria: proteins identified by >1 peptide and that demonstrated a ±>20% change between *APOEx/4* BA41/42 AD and control subjects. The Database for Annotation Visualization and Integrated Discovery (DAVID) was used to test whether synaptic protein sets were enriched in the samples ^51^. To obtain further insight into potential pathways changed in AD synapses, Ingenuity Pathway Analysis (IPA, Ingenuity Systems) was used as previously described ^12, 13, 52^ with the interaction data limited as follows: direct and indirect interactions; experimentally observed data only; 35 molecules per network; 10 networks per dataset. Prediction activation scores (z-scores) were calculated in IPA. Expression clustering was performed in Biolayout Express 3D software by applying Markov clustering algorithms to raw proteomic data (MCL 19 2.2) as previously described in ^13^. All graphs were clustered using Pearson correlation r=0.96.

### Data sharing

Unfiltered proteomics data is included as supplemental table 5. The mass spectrometry proteomics data have also been deposited to the ProteomeXchange Consortium via the PRIDE partner repository ^53^, with the dataset identifier PXD013753. DAVID analysis is provided in supplemental table 2. The full IPA analysis from Figure 3 is in supplemental table 3. Complement proteins identified are in supplemental table 4. Filtered proteomics data used for IPA analysis in figure 3 is provided in supplemental table 6.

## Acknowledgments

Ethics approval and consent to participate: Use of human tissue for post-mortem studies has been reviewed and approved by the Edinburgh Brain Bank ethics committee and the ACCORD medical research ethics committee, AMREC (ACCORD is the Academic and Clinical Central Office for Research and Development, a joint office of the University of Edinburgh and NHS Lothian — Ethics approval reference 15-HV-016). The Edinburgh Brain Bank is a Medical Research Council funded facility with research ethics committee (REC) approval (11/ES/0022). Tissue from 34 donors was used for this study and their details are found in Table 1.

The authors thank our brain tissue donors and their families for their generous donations and Amy Tavendale for her help LC-MS/MS data acquisition. Authors gratefully acknowledge membership of Edinburgh Neuroscience. Some of the control participants in the human study were from the Lothian Birth Cohort 1936, thus we wish to thank the cohort and research team supported by Age UK (Disconnected Mind project) in The University of Edinburgh Centre for Cognitive Ageing and Cognitive Epidemiology, funded by the Biotechnology and Biological Sciences Research Council (BBSRC) and Medical Research Council (MRC) ((MR/K026992/1).

## Consent for publication

Not applicable.

## Authors contributions

Study concept and design: TSJ, TMW, AGH, RH, RJJ; acquisition of data: RH, RJJ, SLE, DL; analysis and interpretation of data: RH, MLH, DK, TMW, TSJ; drafting of the manuscript: RH, TSJ, TMW; critical revision of the manuscript for important intellectual content: TSJ, TMW, SLE, MLE.

## References

1. WHO. Dementia, a global health priority. (ed^(eds). World Health Organisation (2017).

2. Serrano-Pozo A, Frosch MP, Masliah E, Hyman BT. Neuropathological alterations in Alzheimer disease. Cold Spring Harbor perspectives in medicine 1, (2011).

3. Corder EH, et al. Gene dose of apolipoprotein E type 4 allele and the risk of Alzheimer’s disease in late onset families. Science 261, 921–923 (1993).

4. Hudry E, et al. Gene transfer of human Apoe isoforms results in differential modulation of amyloid deposition and neurotoxicity in mouse brain. Science translational medicine 5, 212ra161 (2013).

5. Koffie RM, et al. Apolipoprotein E4 effects in Alzheimer’s disease are mediated by synaptotoxic oligomeric amyloid-beta. Brain 135, 2155–2168 (2012).

6. DeKosky ST, Scheff SW, Styren SD. Structural correlates of cognition in dementia: quantification and assessment of synapse change. Neurodegeneration 5, 417–421 (1996).

7. Terry RD, et al. Physical basis of cognitive alterations in Alzheimer’s disease: synapse loss is the major correlate of cognitive impairment. Ann Neurol 30, 572–580 (1991).

8. Spires-Jones TL, Hyman BT. The intersection of amyloid beta and tau at synapses in Alzheimer’s disease. Neuron 82, 756–771 (2014).

9. Shi Y, et al. ApoE4 markedly exacerbates tau-mediated neurodegeneration in a mouse model of tauopathy. Nature 549, 523–527 (2017).

10. Krasemann S, et al. The TREM2-APOE Pathway Drives the Transcriptional Phenotype of Dysfunctional Microglia in Neurodegenerative Diseases. Immunity 47, 566–581 e569 (2017).

11. Carlyle BC, et al. A multiregional proteomic survey of the postnatal human brain. Nat Neurosci 20, 1787–1795 (2017).

12. Graham LC, et al. Proteomic profiling of neuronal mitochondria reveals modulators of synaptic architecture. Mol Neurodegener 12, 77 (2017).

13. Llavero Hurtado M, et al. Proteomic mapping of differentially vulnerable pre-synaptic populations identifies regulators of neuronal stability in vivo. Sci Rep 7, 12412 (2017).

14. Jones RA, et al. Cellular and Molecular Anatomy of the Human Neuromuscular Junction. Cell Rep 21, 2348–2356 (2017).

15. Bayes A, et al. Human post-mortem synapse proteome integrity screening for proteomic studies of postsynaptic complexes. Mol Brain 7, 88 (2014).

16. Graham LC, et al. Regional molecular mapping of primate synapses during normal healthy ageing. Cell Rep In press, (2019).

17. McGorum BC, et al. Proteomic Profiling of Cranial (Superior) Cervical Ganglia Reveals Beta-Amyloid and Ubiquitin Proteasome System Perturbations in an Equine Multiple System Neuropathy. Mol Cell Proteomics 14, 3072–3086 (2015).

18. Cox J, Mann M. MaxQuant enables high peptide identification rates, individualized p.p.b.-range mass accuracies and proteome-wide protein quantification. Nat Biotechnol 26, 1367–1372 (2008).

19. Cox J, Neuhauser N, Michalski A, Scheltema RA, Olsen JV, Mann M. Andromeda: a peptide search engine integrated into the MaxQuant environment. J Proteome Res 10, 1794–1805 (2011).

20. Pickett EK, et al. Region-specific depletion of synaptic mitochondria in the brains of patients with Alzheimer’s disease. Acta Neuropathol 136, 747–757 (2018).

21. Kay KR, et al. Studying synapses in human brain with array tomography and electron microscopy. Nat Protoc 8, 1366–1380 (2013).

22. Henstridge CM, Spires-Jones T. Beyond the neuron - Cellular interactions early in Alzheimer’s disease pathogenesis. Nature Reviews Neuroscience in press, (2019).

23. Izzo NJ, et al. Alzheimer’s therapeutics targeting amyloid beta 1-42 oligomers II: Sigma-2/PGRMC1 receptors mediate Abeta 42 oligomer binding and synaptotoxicity. PLoS One 9, e111899 (2014).

24. Hong S, et al. Complement and microglia mediate early synapse loss in Alzheimer mouse models. Science, (2016).

25. Shi Q, et al. Complement C3 deficiency protects against neurodegeneration in aged plaque-rich APP/PS1 mice. Science translational medicine 9, (2017).

26. Dejanovic B, et al. Changes in the Synaptic Proteome in Tauopathy and Rescue of Tau-Induced Synapse Loss by C1q Antibodies. Neuron, (2018).

27. Litvinchuk A, et al. Complement C3aR Inactivation Attenuates Tau Pathology and Reverses an Immune Network Deregulated in Tauopathy Models and Alzheimer’s Disease. Neuron, (2018).

28. Riad A, et al. Sigma-2 Receptor/TMEM97 and PGRMC-1 Increase the Rate of Internalization of LDL by LDL Receptor through the Formation of a Ternary Complex. Sci Rep 8, 16845 (2018).

29. Eaton SL, et al. Total protein analysis as a reliable loading control for quantitative fluorescent Western blotting. PLoS One 8, e72457 (2013).

30. Andreev VP, et al. Label-free quantitative LC-MS proteomics of Alzheimer’s disease and normally aged human brains. J Proteome Res 11, 3053–3067 (2012).

31. Musunuri S, et al. Quantification of the brain proteome in Alzheimer’s disease using multiplexed mass spectrometry. J Proteome Res 13, 2056–2068 (2014).

32. Zhang Q, Ma C, Gearing M, Wang PG, Chin LS, Li L. Integrated proteomics and network analysis identifies protein hubs and network alterations in Alzheimer’s disease. Acta Neuropathol Commun 6, 19 (2018).

33. Wang S, et al. Quantitative proteomics identifies altered O-GlcNAcylation of structural, synaptic and memory-associated proteins in Alzheimer’s disease. J Pathol 243, 78–88 (2017).

34. Reinders NR, et al. Amyloid-beta effects on synapses and memory require AMPA receptor subunit GluA3. Proc Natl Acad Sci U S A 113, E6526–E6534 (2016).

35. Baglietto-Vargas D, et al. Impaired AMPA signaling and cytoskeletal alterations induce early synaptic dysfunction in a mouse model of Alzheimer’s disease. Aging Cell, e12791 (2018).

36. Duits FH, et al. Synaptic proteins in CSF as potential novel biomarkers for prognosis in prodromal Alzheimer’s disease. Alzheimers Res Ther 10, 5 (2018).

37. Portelius E, et al. Cerebrospinal fluid neurogranin concentration in neurodegeneration: relation to clinical phenotypes and neuropathology. Acta Neuropathol 136, 363–376 (2018).

38. Kvartsberg H, et al. Cerebrospinal fluid levels of the synaptic protein neurogranin correlates with cognitive decline in prodromal Alzheimer’s disease. Alzheimers Dement 11, 1180–1190 (2015).

39. Lleó A, et al. Changes in Synaptic Proteins Precede Neurodegeneration Markers in Preclinical Alzheimer’s Disease Cerebrospinal Fluid. Molecular &amp; Cellular Proteomics 18, 546–560 (2019).

40. Lin YT, et al. APOE4 Causes Widespread Molecular and Cellular Alterations Associated with Alzheimer’s Disease Phenotypes in Human iPSC-Derived Brain Cell Types. Neuron 98, 1294 (2018).

41. Atagi Y, et al. Apolipoprotein E Is a Ligand for Triggering Receptor Expressed on Myeloid Cells 2 (TREM2). J Biol Chem 290, 26043–26050 (2015).

42. Yin C, et al. ApoE attenuates unresolvable inflammation by complex formation with activated C1q. Nat Med, (2019).

43. Lasagna-Reeves CA, Castillo-Carranza DL, Sengupta U, Clos AL, Jackson GR, Kayed R. Tau oligomers impair memory and induce synaptic and mitochondrial dysfunction in wild-type mice. Mol Neurodegener 6, 39 (2011).

44. Manczak M, Kandimalla R, Fry D, Sesaki H, Reddy PH. Protective effects of reduced dynamin-related protein 1 against amyloid beta-induced mitochondrial dysfunction and synaptic damage in Alzheimer’s disease. Hum Mol Genet 25, 5148–5166 (2016).

45. Reddy PH, Beal MF. Amyloid beta, mitochondrial dysfunction and synaptic damage: implications for cognitive decline in aging and Alzheimer’s disease. Trends Mol Med 14, 45–53 (2008).

46. Chang S, ran Ma T, Miranda RD, Balestra ME, Mahley RW, Huang Y. Lipid- and receptor-binding regions of apolipoprotein E4 fragments act in concert to cause mitochondrial dysfunction and neurotoxicity. Proc Natl Acad Sci U S A 102, 18694–18699 (2005).

47. Henstridge CM, et al. Synapse loss in the prefrontal cortex is associated with cognitive decline in amyotrophic lateral sclerosis. Acta Neuropathol 135, 213–226 (2018).

48. Tai HC, Serrano-Pozo A, Hashimoto T, Frosch MP, Spires-Jones TL, Hyman BT. The synaptic accumulation of hyperphosphorylated tau oligomers in Alzheimer disease is associated with dysfunction of the ubiquitin-proteasome system. Am J Pathol 181, 1426–1435 (2012).

49. Hesse R, et al. sAPPβ and sAPPα increase structural complexity and E/I input ratio in primary hippocampal neurons and alter Ca. Exp Neurol 304, 1–13 (2018).

50. Wisniewski JR, Zougman A, Nagaraj N, Mann M. Universal sample preparation method for proteome analysis. Nat Methods 6, 359–362 (2009).

51. Huang DW, Sherman BT, Lempicki RA. Systematic and integrative analysis of large gene lists using DAVID bioinformatics resources. Nat Protoc 4, 44–57 (2009).

52. Wishart TM, et al. Differential proteomics analysis of synaptic proteins identifies potential cellular targets and protein mediators of synaptic neuroprotection conferred by the slow Wallerian degeneration (Wlds) gene. Mol Cell Proteomics 6, 1318–1330 (2007).

53. Perez-Riverol Y, et al. The PRIDE database and related tools and resources in 2019: improving support for quantification data. Nucleic Acids Res 47, D442–D450 (2019).

