## Supplementary material for "Comparative profiling of the synaptic proteome from Alzheimer’s disease patients with focus on the APOE genotype": Supplemebtal table 1

Supplemental Table 1: Previous proteomics studies of post mortem human AD brain.

| **Title** | **Year** | **Type of Proteomics** | **Brain area and subcellular fraction used** | **# Changes Detected** | **Major Conclusions** |
| --- | --- | --- | --- | --- | --- |
| Proteomic analysis of the brain in Alzheimer's disease: molecular phenotype of a complex disease process  (Schonberger, Edgar et al. 2001) | 2001 | 2D electrophoresis followed by in gel trypsin digestion and then HPLC and sequence identification | Whole tissue homogenate from Hippocampus (Hp), temporal cortex (tCx), entorhinal cortex (EC), Cerebellum (Cb), cingulate gyrus (cGy) and sensorimotor cortex (sCx) from AD and matched controls | 76 in Hp  62 in tCx  39 in EC  34 in Cb  125 in cGy  75 in sCx | Protein differences in AD are not specific to regions of severe degeneration. Proteins changed in AD were involved in synaptic neurotransmission, stress response, lipid transport, glycolysis, and known Diabetes pathways |
| Proteomic Profiling and Neurodegeneration in Alzheimer’s Disease  (Tsuji, Shiozaki et al. 2002) | 2002 | 2D electrophoresis, followed by in gel digestion and LC-MS/MS | Whole tissue homogenate of temporal cortex from AD and controls | 35 | This seems to have been a proof of concept for the techniques used. |
| Proteomics Analysis of the Alzheimer’s Disease Hippocampal Proteome  (Sultana, Boyd-Kimball et al. 2007) | 2007 | 2D electrophoresis, followed by in gel digestion and MALDITOF mass spectrometry | AD and control inferior parietal lobule and hippocampus | 18 | Found changes in energy related enzymes, scaffolding proteins particularly HSP70, structural proteins, cell cycle, tau phosphorylation and Aβ production |
| An Increase in S-Glutathionylated Proteins in the Alzheimer’s Disease Inferior Parietal Lobule, a Proteomics Approach  (Newman, Sultana et al. 2007) | 2007 | redox proteomics: 2D electrophoresis | AD and control inferior parietal lobule and hippocampus | 4 | specific proteins have an increased S-glutathionylation in the AD brain which probably diminishes their activity |
| Analysis of microdissected neurons by 18O mass spectrometry reveals altered protein expression in Alzheimer’s disease  (Koffie, Hashimoto et al. 2012) | 2012 | AD samples were labelled with O18 while control samples were not. Samples were then trypsinized and LC-MS/MS was used | Microdissected neurons from the hippocampus of AD and control tissue | 68 | Many of the proteins found to be different were involved in glycolysis |
| Analysis of a membrane-enriched proteome from postmortem human brain tissue in Alzheimer's disease.  (Donovan, Higginbotham et al. 2012) | 2012 | Trypsin digest followed by reverse phase LC-MS/MS | Membrane enriched sample from frontal cortex of AD v matched controls | 13 | Tau was the most significantly changed protein in AD cases |
| Label-Free Quantitative LC-MS Proteomics of Alzheimer’s Disease and Normally Aged Human Brains  (Andreev, Petyuk et al. 2012) | 2012 | Accurate mass and time tag with a LTQ Orbitrap mass spec | Temporal lobe homogenate | 197 | Protein families shown to changed included signal transduction, regulation of protein phosphorylation, immune response, cytoskeleton organization, lipid metabolism, energy production, and cell death. Also highlighted are the proteins that differed from published literature |
| **Title** | **Year** | **Type of Proteomics** | **Brain area and subcellular fraction used** | **# Changes Detected** | **Major Conclusions** |
| Proteomic Analysis of Postsynaptic Density in Alzheimer Disease  (Zhou, Jones et al. 2013) | 2013 | Label free LC-MS/MS | PSD enriched from cortex of AD, probable AD and control cases using a sucrose gradient | 25 | The family of proteins that regulate actin dynamics were changed. |
| Brain site-specific proteome changes in aging-related dementia  (Manavalan, Mishra et al. 2013) | 2013 | Proteins were digested in gel following 2D electrophoresis then iTRAQ tags were added and LC-MS/MS was used. | Whole tissue homogenate from the hippocampus (Hp), parietal cortex (pCx) and cerebellum (Cb) of AD and age matched control females | 31 Total  22 in Hp  8 in pCx  16 in Cb | Different areas of the brain have different and overlapping proteins changes in AD. Tau was found to be different in AD in both the Hp and the Cb and Aβ was different in both the Hp and pCx. |
| Semiquantitative proteomic analysis of human hippocampal tissues from Alzheimer's disease and age-matched control brains  (Begcevic, Kosanam et al. 2013) | 2013 | Trypsin digestion followed by semiquantitative label-free LC-MS/MS | Whole tissue homogenate from AD and control hippocampus | 204 detected only in AD and 600 in only control | Many of the proteins of the AD Hp proteome are involved in protein binding, catalytic activity and nucleotide binding. 40 of the AD specific and 106 of the control specific proteins are detected in CSF indicating they could be biomarkers |
| The synaptic proteome in Alzheimer’s disease  (Chang, Nouwens et al. 2013) | 2013 | 2D differential in-gel electrophoresis, then time of flight mass spec on dots of interest. | Synaptosomes of 2 affected areas- hippocampus and inferior temporal gyrus and 2 unaffected areas- occiptial cortex and motor cortex. | 26 | No significant difference between the unaffected areas in both AD and non AD cases therefore these were used to normalize expression in affected areas. |
| An investigation of the molecular mechanisms engaged before  and after the development of Alzheimer disease neuropathology  in Down syndrome: a proteomics approach  (Cenini, Fiorini et al. 2014) | 2014 | 2D differential in-gel electrophoresis followed by in gel digestion and then mass spectrometry | Frontal cortex homogenate from AD, down syndrome (DS), DS with AD tissue (AD/DS) and both young (YC) and old controls (OC) | 7 (DS v YC)  3 (AD/DS v OC)  3 (DS v AD/DS)  10 (YC v OC) | Redox protein changes are important in DS and AD. ApoE is found to be less abundant in young DS brains compared with young controls. |
| Quantification of the brain proteome in Alzheimer's disease using multiplexed mass spectrometry.  (Musunuri, Wetterhall et al. 2014) | 2014 | Trypsin digestion followed by stable-isotope dimethyl labelling and then nanoLC-MS/MS | Whole tissue homogenate of temporal neocortex from AD and matched controls | 69 | Proteins increased in AD were involved in metabolic processes, oxidative stress and inflammation, proteins decreased in AD were involved in altered synaptic function and signal transduction |
| Differential expression of proteins in brain regions of Alzheimer's disease patients  (Zahid, Oellerich et al. 2014) | 2014 | 2D electrophoresis, followed by in gel digestion and ESI-QTOF-MS/MS | Whole tissue homogenate of Hippocampus (Hp), substantia nigra (SN) and frontal cortex (fCx) from AD and control cases | 48 total  13 in Hp  20 in SN  22 in fCx | Protein changes were differently regulated in the different brain areas. Differentially expressed proteins from AD Hp, fCx and SN are involved in metabolism, transport and the cytoskeleton. |
| **Title** | **Year** | **Type of Proteomics** | **Brain area and subcellular fraction used** | **# Changes Detected** | **Major Conclusions** |
| Apolipoprotein E*4 (APOE*4) Genotype Is Associated with Altered Levels of Glutamate Signaling Proteins and Synaptic Coexpression Networks in the Prefrontal Cortex in Mild to Moderate Alzheimer Disease (Sweet, MacDonald et al. 2016) | 2016 | Trypsin digestion followed by C13 labeling and LC-MS/MS | Synaptosomes from AD, FTD, and control humans as well as ApoE3 and ApoE4 mice were isolated using a sucrose gradient from the Dorsolateral prefrontal cortex (DLPC) and the Entorhinal cortex (EC) | 1 in DLPC in AD  95 in DLPC in FTD  95 in EC in AD | The AD human samples clustered into two groups which were enriched for ApoE3 or ApoE4 genotypes. The main pathway that differed between the two groups were glutamate signalling indicating that ApoE4 could play a role in reducing glutamate signalling although this finding was not recapitulated in the ApoE4 mice. |
| Olfactory bulb neuroproteomics reveals a chronological perturbation of survival routes and a disruption of prohibitin complex during Alzheimer's disease progression (Lachen-Montes et al. 2017) | 2017 | Trypsin digestion, label free LC-MS/MS | Whole tissue homogenate of Olfactory bulb | 1311 IDs, 278 changed in AD | stage-dependent synaptic proteostasis impairment, mitochondrial dysfunction, modulation of tau and APP interactomes in AD |
| Integrated proteomics and network analysis identifies protein hubs and network alterations in Alzheimer's disease (Zhang et al 2018) | 2018 | Trypsin digestion, label free LC-MS/MS | Whole tissue homogenate from frontal cortex of AD and controls | 6,679 IDs, 1,968 anaysed, 487 changed in AD | Observed altered proteostasis, RNA homeostasis, immune response, neuroinflammation, synaptic transmission, vesicular transport, cell signaling, cellular metabolism, lipid homeostasis, mitochondrial dynamics and function, cytoskeleton organization, and myelin-axon interactions. No correlation with APOE genotype in their study. |
| Deep proteomic network analysis of Alzheimer's disease brain reveals alterations in RNA binding proteins and RNA splicing associated with disease (Johnsson et al. 2018) | 2018 | TMT labelling, LC-MS/MS | Whole tissue homogenate from frontal cortex (BA9) of AD, asymptomatic AD, and controls | 6,533 IDs, 350 changed in AD vs asymptAD | RNA binding proteins were altered in AD and in AD there was alternative splicing including of risk genes. |
| Global quantitative analysis of the human brain proteome in Alzheimer's and Parkinson's Disease (Ping et al. 2018) | 2018 | Trypsin digestion, TMT labelling, synchronous precursor selection-based MS3 LC-MS/MS | Whole tissue homogenate from frotnal (BA9) and cingulate (BA24) cortex of AD, PD, mixed AD/PD and controls | >10,000 IDs | Strong linear correlation between protein abundance in different brain regions. Disease effects were not analysed. |

**Search terms:** ("proteomics"[MeSH Terms] OR "proteomics"[Title/Abstract]) AND ("alzheimer disease"[MeSH Terms] OR ("alzheimer"[ Title/Abstract] AND "disease"[ Title/Abstract]) OR "alzheimer disease"[ Title/Abstract] OR "alzheimer"[ Title/Abstract]) AND ("brain"[MeSH Terms] OR "brain"[ Title/Abstract]) AND ("humans"[MeSH Terms] OR "humans"[ Title/Abstract] OR "human"[ Title/Abstract])

**Pubmed hits:** 211 before filtering, 19 after filtering

**Inclusion criteria:** (1) containing original proteomic data (2) from human Alzheimer and control brain tissue samples.

**Exclusion criteria:** analysis of (1) isolated protein aggregates, (1) insoluble fractions, or (3) IP of specific protein

**References**

Andreev, V. P., V. A. Petyuk, H. M. Brewer, Y. V. Karpievitch, F. Xie, J. Clarke, D. Camp, R. D. Smith, A. P. Lieberman, R. L. Albin, Z. Nawaz, J. El Hokayem and A. J. Myers (2012). "Label-free quantitative LC-MS proteomics of Alzheimer's disease and normally aged human brains." J Proteome Res **11**(6): 3053-3067.

Begcevic, I., H. Kosanam, E. Martinez-Morillo, A. Dimitromanolakis, P. Diamandis, U. Kuzmanov, L. N. Hazrati and E. P. Diamandis (2013). "Semiquantitative proteomic analysis of human hippocampal tissues from Alzheimer's disease and age-matched control brains." Clin Proteomics **10**(1): 5.

Cenini, G., A. Fiorini, R. Sultana, M. Perluigi, J. Cai, J. B. Klein, E. Head and D. A. Butterfield (2014). "An investigation of the molecular mechanisms engaged before and after the development of Alzheimer disease neuropathology in Down syndrome: a proteomics approach." Free Radic Biol Med **76**: 89-95.

Chang, R. Y., A. S. Nouwens, P. R. Dodd and N. Etheridge (2013). "The synaptic proteome in Alzheimer's disease." Alzheimers Dement **9**(5): 499-511.

Donovan, L. E., L. Higginbotham, E. B. Dammer, M. Gearing, H. D. Rees, Q. Xia, D. M. Duong, N. T. Seyfried, J. J. Lah and A. I. Levey (2012). "Analysis of a membrane-enriched proteome from postmortem human brain tissue in Alzheimer's disease." Proteomics Clin Appl **6**(3-4): 201-211.

Johnson, E. C. B., E. B. Dammer, D. M. Duong, L. Yin, M. Thambisetty, J. C. Troncoso, J. J. Lah, A. I. Levey and N. T. Seyfried (2018). "Deep proteomic network analysis of Alzheimer's disease brain reveals alterations in RNA binding proteins and RNA splicing associated with disease." Mol Neurodegener **13**(1): 52.

Koffie, R. M., T. Hashimoto, H. C. Tai, K. R. Kay, A. Serrano-Pozo, D. Joyner, S. Hou, K. J. Kopeikina, M. P. Frosch, V. M. Lee, D. M. Holtzman, B. T. Hyman and T. L. Spires-Jones (2012). "Apolipoprotein E4 effects in Alzheimer's disease are mediated by synaptotoxic oligomeric amyloid-beta." Brain **135**(Pt 7): 2155-2168.

Lachen-Montes, M., A. Gonzalez-Morales, M. V. Zelaya, E. Perez-Valderrama, K. Ausin, I. Ferrer, J. Fernandez-Irigoyen and E. Santamaria (2017). "Olfactory bulb neuroproteomics reveals a chronological perturbation of survival routes and a disruption of prohibitin complex during Alzheimer's disease progression." Sci Rep **7**(1): 9115.

Manavalan, A., M. Mishra, L. Feng, S. K. Sze, H. Akatsu and K. Heese (2013). "Brain site-specific proteome changes in aging-related dementia." Exp Mol Med **45**: e39.

Musunuri, S., M. Wetterhall, M. Ingelsson, L. Lannfelt, K. Artemenko, J. Bergquist, K. Kultima and G. Shevchenko (2014). "Quantification of the brain proteome in Alzheimer's disease using multiplexed mass spectrometry." J Proteome Res **13**(4): 2056-2068.

Newman, S. F., R. Sultana, M. Perluigi, R. Coccia, J. Cai, W. M. Pierce, J. B. Klein, D. M. Turner and D. A. Butterfield (2007). "An increase in S-glutathionylated proteins in the Alzheimer's disease inferior parietal lobule, a proteomics approach." J Neurosci Res **85**(7): 1506-1514.

Ping, L., D. M. Duong, L. Yin, M. Gearing, J. J. Lah, A. I. Levey and N. T. Seyfried (2018). "Global quantitative analysis of the human brain proteome in Alzheimer's and Parkinson's Disease." Sci Data **5**: 180036.

Schonberger, S. J., P. F. Edgar, R. Kydd, R. L. Faull and G. J. Cooper (2001). "Proteomic analysis of the brain in Alzheimer's disease: molecular phenotype of a complex disease process." Proteomics **1**(12): 1519-1528.

Sultana, R., D. Boyd-Kimball, J. Cai, W. M. Pierce, J. B. Klein, M. Merchant and D. A. Butterfield (2007). "Proteomics analysis of the Alzheimer's disease hippocampal proteome." J Alzheimers Dis **11**(2): 153-164.

Sweet, R. A., M. L. MacDonald, C. M. Kirkwood, Y. Ding, T. Schempf, J. Jones-Laughner, J. Kofler, M. D. Ikonomovic, O. L. Lopez, M. E. Garver, N. F. Fitz, R. Koldamova and N. A. Yates (2016). "Apolipoprotein E*4 (APOE*4) Genotype Is Associated with Altered Levels of Glutamate Signaling Proteins and Synaptic Coexpression Networks in the Prefrontal Cortex in Mild to Moderate Alzheimer Disease." Mol Cell Proteomics **15**(7): 2252-2262.

Tsuji, T., A. Shiozaki, R. Kohno, K. Yoshizato and S. Shimohama (2002). "Proteomic profiling and neurodegeneration in Alzheimer's disease." Neurochem Res **27**(10): 1245-1253.

Zahid, S., M. Oellerich, A. R. Asif and N. Ahmed (2014). "Differential expression of proteins in brain regions of Alzheimer's disease patients." Neurochem Res **39**(1): 208-215.

Zhang, Q., C. Ma, M. Gearing, P. G. Wang, L. S. Chin and L. Li (2018). "Integrated proteomics and network analysis identifies protein hubs and network alterations in Alzheimer's disease." Acta Neuropathol Commun **6**(1): 19.

Zhou, J., D. R. Jones, D. M. Duong, A. I. Levey, J. J. Lah and J. Peng (2013). "Proteomic analysis of postsynaptic density in Alzheimer's disease." Clin Chim Acta **420**: 62-68.
